# Metabolic Regulation of Mitochondrial Morphologies in Pancreatic Beta Cells: Bioenergetics-Mitochondrial Dynamics Coupling

**DOI:** 10.1101/2021.10.04.462897

**Authors:** Wen-Wei Tseng, Ching-Hsiang Chu, Chen Chang, Yi-Ju Lee, Shirui Zhao, Yi-Ping Ho, An-Chi Wei

## Abstract

Cellular bioenergetics and mitochondrial dynamics are crucial for the secretion of insulin by pancreatic beta cells in response to elevated blood glucose concentrations. To obtain better insights into the interactions between energy production and mitochondrial fission/fusion dynamics, we combine live-cell mitochondria imaging with biophysical-based modeling and network analysis to elucidate the principle regulating mitochondrial morphology to match metabolic demand in pancreatic beta cells. A minimalistic differential equation-based model for beta cells was constructed to include glycolysis, oxidative phosphorylation, simple calcium dynamics, and graph-based fission/fusion dynamics controlled by ATP synthase flux and proton leak flux. The model revealed that mitochondrial fission occurs in response to hyperglycemia, starvation, ATP synthase inhibition, uncoupling, and diabetic condition, in which the rate of proton leak exceeds the rate of mitochondrial ATP synthesis. Under these metabolic challenges, the propensities of tip-to-tip fusion events simulated from the microscopic images of the mitochondrial networks were lower than those in the control group and prevented mitochondrial network formation. The modeling and network analysis could serve as the basis for further detailed research on the mechanisms of bioenergetics and mitochondrial dynamics coupling.

## Introduction

Mitochondria, in charge of the generation of adenosine triphosphate (ATP) via oxidative phosphorylation (OXPHOS), are motile organelles exhibiting dynamic structures owing to the fission and fusion cycles.[1] Fission-fusion cycles, which are typically referred to as mitochondrial dynamics, are related to the regulation of energy production and the quality control of the mitochondrial network.[2,3] Mitochondrial fusion and biogenesis maintain the mitochondrial mass and network [4]. Moreover, quality control is achieved by asymmetric fission, which splits one mitochondrion into two daughter mitochondria with different mitochondrial membrane potentials. Deenergized mitochondria are subsequently removed through mitophagy. At the molecular level, fusion and fission events are driven by large GTPases. Mitofusin 1 (Mfn1) and 2 (Mfn2) on the outer mitochondrial membrane (OMM) and optic atrophy 1 (OPA1) on the inner mitochondrial membrane (IMM) are in charge of fusion events, whereas dynamin-related/-like protein 1 (Drp1) and Dynamin2 (Dnm2) are responsible for fission events.[1]

The mitochondrial network may shift to different morphologies depending on the metabolic task. Recent studies have linked the nutrient supply and energy demand to mitochondrial dynamics, which implies that the mitochondrial architecture adapts to metabolic demands [5,6]. Taking rat insulinoma (INS) cells as an example [3], mitochondrial networks are more tubular during the G1 to S phases for energy production and biosynthesis, whereas mitochondria are more fragmented during the G2 to M phases to prepare for an equal distribution of the mitochondrion into the daughter cells. In cancer cells, mitochondrial dynamics are also crucial for their metabolic rewiring, metastasis abilities, drug resistance, and cancer stem cell survival.[7–10] Pancreatic beta cells sense the glucose concentration by coupling glycolysis to the citric acid cycle (CAC) and OXPHOS to synthesize ATP, and this step increases the ATP-to-ADP ratio, closes ATP-inhibited potassium channels (KATP channels), triggers calcium influx, and excretes insulin vesicles.[11,12] The process, which is called glucose-stimulated insulin secretion (GSIS), is correlated with mitochondrial morphology changes.[13–15] Glucose stimulation could induce short-term (approximately 1 hour) mitochondrial fragmentation and recovery.[14]

In contrast, perturbations in mitochondrial dynamics are associated with the deterioration of mitochondrial network quality, mitochondrial dysfunction, decreased ATP synthesis capacity, impaired calcium homeostasis, and even cell death.[16] Insulin resistance and type 2 diabetes (T2DM) are associated with hampered mitochondrial functions in OXPHOS and citric acid cycle (CAC) metabolism [17,18], which results in attenuation of the sensitivity to glucose stimulation of beta cells to secrete insulin. Fragmented mitochondrial network morphologies have also been found in diabetic beta cells [10,19–23]. The artificial blockage of fission proteins in INS-1E cells hinders glucose sensing and insulin secretion.[14]

A variety of studies have evaluated the mitochondrial morphology and mitochondrial network fission/fusion dynamics. For instance, fluorescence microscopy provides direct evidence of the mitochondrial network shapes and measurements of their motility, fission, and fusion rates.[24–26] Image preprocessing and analysis techniques are indispensable for revealing information from these data and further providing quantitative evidence of the results and discoveries based on mitochondrial dynamics. [27–29]

In contrast, *in silico* models describe the relationships between substrate input and mitochondrial ATP production using computer simulations. The widely adopted [30–35] mathematical model of beta-cell mitochondria by Magnus and Kaiser [36] described the influence of intracellular calcium dynamics and adenylate levels on ATP synthesis and the mitochondrial membrane potential. The model was further simplified [37] and then revised [38] to reveal the frequency of the response of ATP synthesis to respiring substrates and the cytosolic concentrations of calcium. The INS beta-cell models constructed by Fridlyand et al.[39–42] are focused on ATP production, the plasma membrane potential dynamics, intracellular calcium oscillations, signal transduction, and insulin secretion in response to glucose stimulation, i.e., glucose-stimulated insulin secretion (GSIS). Although the mitochondrial bioenergetics models describe ATP synthesis upon substrate addition, the mitochondria are regarded as a single entity without fission/fusion dynamics.

Alternatively, mitochondria could be treated as individual agents undergoing fission/fusion dynamics, and one could evaluate the overall mitochondrial network mass, structure, and quality [43–45]. Patel and coworkers [16] devised an agent-based model (ABM) to simulate mitochondrial movement, fission, fusion, mitophagy, and biogenesis processes and revealed that the selective mitophagy of damaged mitochondria improves the overall mitochondrial health. Sukhorukov and coworkers [46] described mitochondrial fission-fusion dynamics as dissociation and association of nodes to simulate the observed fluorescence microscopic images of mitochondrial networks in HeLa cells. Their idea was further expanded by Shah and coworkers [20] to estimate the fission/fusion rate differences in healthy and unhealthy cells.

However, only a few studies have combined bioenergetics and mitochondrial fission-fusion dynamics due to the complexity involved in the mathematical model and the scarcity of data related to fusion and fission rates from microscopic observations obtained under the microscope. One of the few examples is Kornick’s population-based model [47], which is a minimalistic ODE-based model simulating the dynamics of fragmented/fused and healthy/unhealthy mitochondrial populations and is dependent on the generation and consumption of ATP. However, the mathematical descriptors for bioenergetics and fission/fusion rates were set based on constant ratios of healthy compared with unhealthy mitochondria without considering any mechanistic details of OXPHOS and the associated effects on mitochondrial dynamics.

To this end, our study was set to bridge among cellular bioenergetics, mitochondrial network dynamics (as illustrated in Fig. 1), and microscopic observations together. We simulated the model and matched the mitochondrial network pattern changes observed by microscopy under various metabolic conditions, such as changing the glucose concentration and adding chemicals that affect the mitochondrial respiratory chain, to gain better insight into the bioenergetic coupling of mitochondrial dynamics in pancreatic beta cells. The model simulations reveal mechanisms for mitochondrial dynamics regulation that link the nutrient environment to mitochondrial dynamics and bioenergetics and are relate to progressive mitochondrial dysfunction in metabolic diseases.

**Fig. 1:**
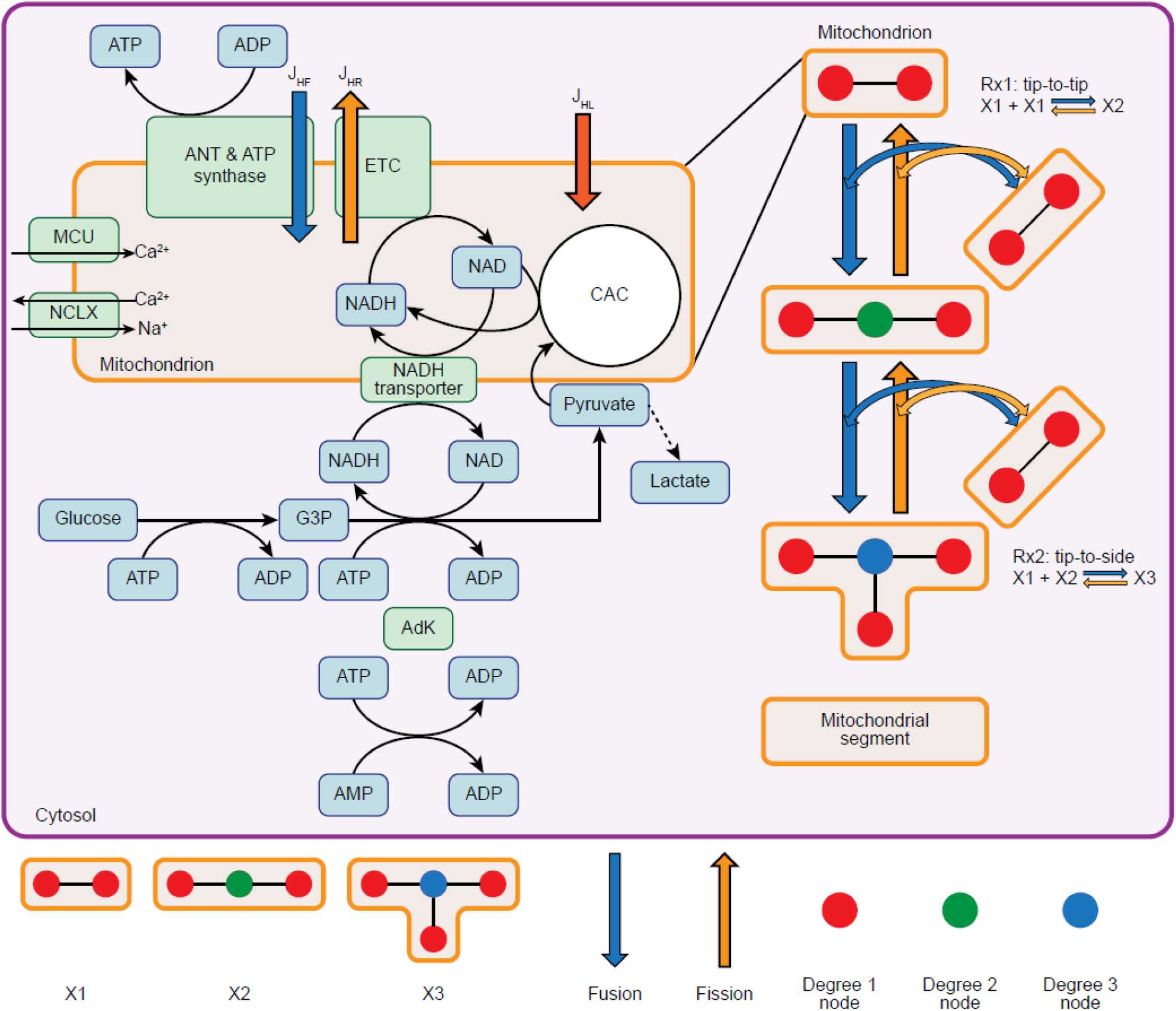
Schematics of the mitochondrial bioenergetics and dynamics model. The *in silico* model for mitochondrial bioenergetics and dynamics is built upon a coupled system of ordinary differential equations (ODEs). This model includes the bioenergetic and fission-fusion dynamics responses to glucose stimulation in pancreatic cells. The left half represents glucose-stimulated insulin secretion (GSIS), which involves the coupling of glycolysis in the cytosol and OXPHOS in the mitochondria, and a minimalistic model of calcium fluxes between the cytosol and mitochondria. In contrast, the right half represents the fission-fusion dynamics of the mitochondrial network. The relative fission-fusion rate is determined by the ATP synthase rate and the proton leak rate. ANT: adenine nucleotide translocator. MCU: calcium uniporter. NCLX: mitochondrial sodium-calcium exchanger (sodium-calcium-lithium exchanger, NCLX). ETC: electron transport chain, including complexes I, II, III, and IV. CAC: citric acid cycle. G3P: glyceraldehyde-3-phosphate. NAD: nicotinamide adenine dinucleotide. NADH: reduced nicotinamide adenine dinucleotide. J_HR_: flux of proton pumping of ETC. JHL: proton leak flux. JHF: proton flux through ATP synthase (F1-Fo ATPase). The mathematical details of the ODE model can be found in the Materials and Methods section and the Supplementary Materials.

## Results

### Glucose stimulates mitochondrial bioenergetics with morphological changes

Glucose is a stimulus signal for metabolic-secretion coupling. Mitochondria in pancreatic beta-cells respond to increases in the blood glucose, which leads to an increased ATP/ADP ratio by increasing oxidative metabolism and increasing the intracellular calcium concentrations to trigger insulin granule exocytosis. Exploration of the mechanism through which the glucose concentration affects mitochondrial dynamics is the first step to understanding how the cellular metabolism regulates mitochondrial dynamics. High glucose concentration increased the steady-state levels of glycolysis metabolites (G3P, pyruvate), the CAC product (NADH), and the OXPHOS products (ATP and ATP-to-ADP ratio) (Fig. 2 (A-G)). Glycolytic flux through glucokinase (GK) drove the downstream reactions of CAC and OXPHOS. Increased cytosolic calcium from the increased ATP-to-ADP ratio resulted in calcium influx into the mitochondria through the mitochondrial calcium uniporter (MCU). Increases in the mitochondrial calcium further enhanced the reactions of CAC and OXPHOS to produce even more ATP.

**Fig. 2:**
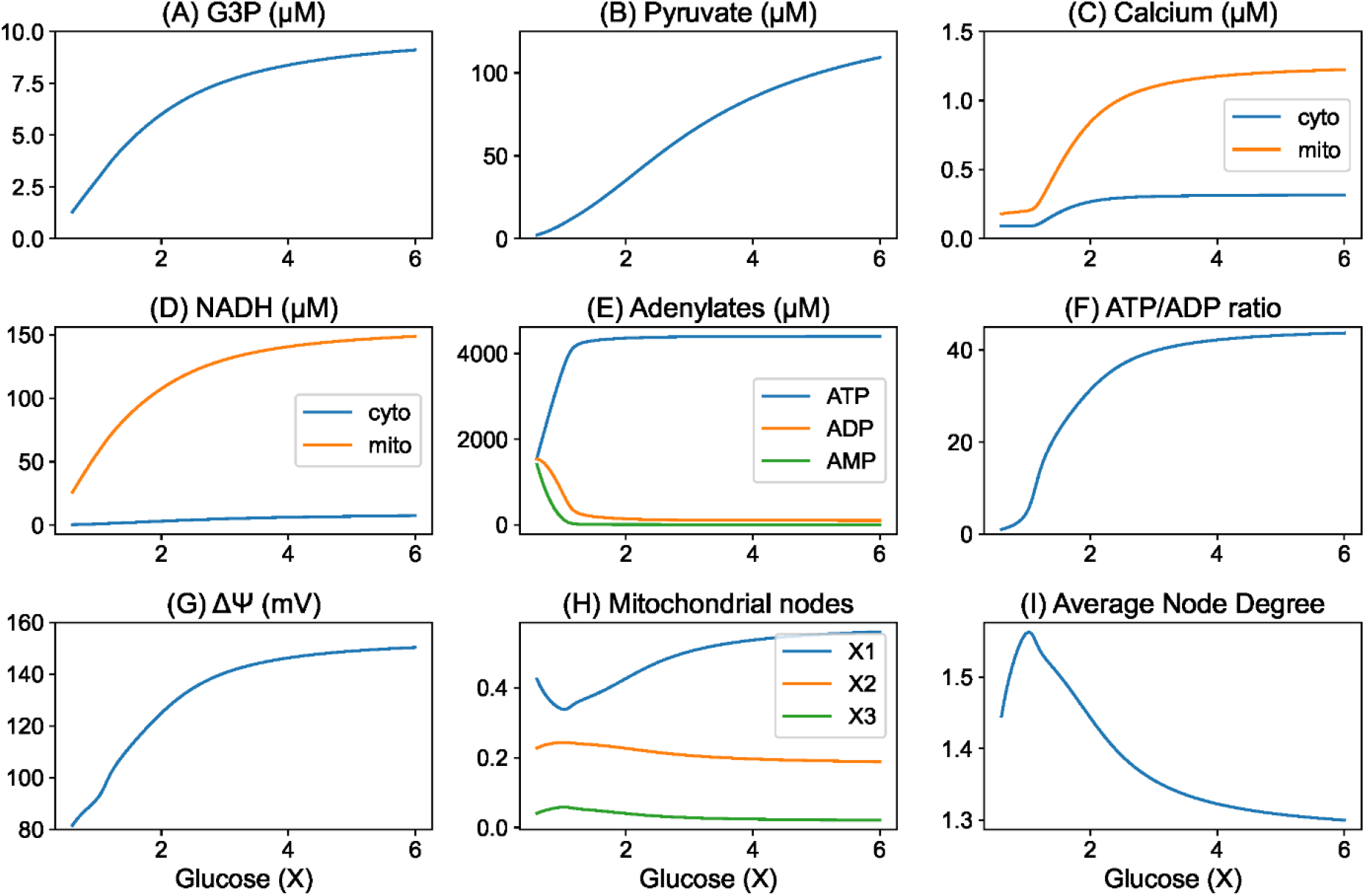
Steady states obtained with a range of glucose concentrations. Steady-state values of (A) G3P (in μM), (B) pyruvate (in μM), (C) calcium (in μM), (D) NADH (in μM), (E) ATP, ADP, and AMP (in μM), (F) the ATP/ADP ratio, (G) the mitochondrial membrane potential (in millivolts, mV), (H) the mitochondrial population, and (I) the average degree of nodes were compared under different glucose concentrations. The x-axis represents relative glucose concentrations, and 5 mM is denoted as 1X.

Moreover, the mitochondrial network became more fragmented in response to elevated glucose concentrations. The population of terminal degree-1 nodes (Fig. 2 (H)) increased, and the average degree of the nodes decreased (Fig. 2 (I)). The mitochondrial network showed the highest degree of fusion with the highest average degree of nodes at a resting glucose concentration of 1X.

### Interactions between mitochondrial energetics and dynamics

To further dissect the relationship between mitochondrial bioenergetics and mitochondrial dynamics, we orthogonally varied both the glucose concentration and one of the mitochondrial parameters: ATP synthase, ETC, and proton leakage activities. We then collected the steady states of the average degree of nodes, the mitochondrial membrane potential, and the ATP/ADP ratio. In all scenarios, when the glucose levels increased, the decreased average degree of nodes showed a tendency of mitochondrial network fragmentation, and both the ATP/ADP ratio and mitochondrial membrane potential increased (Fig. 3). The inhibition of ATP synthase activity resulted in higher mitochondrial membrane potentials, a lower ATP/ADP ratio, and a more fragmented mitochondrial network. In contrast, the inhibition of electron transport chain (ETC) activity moderately suppressed the mitochondrial membrane potential and the ATP/ADP ratio and generated a fused mitochondrial network. An increase in proton leakage dissipated the mitochondrial membrane potential, decreased the ATP/ADP ratio, and scattered the mitochondrial network (Fig. 3 and Table 1).

**Fig. 3:**
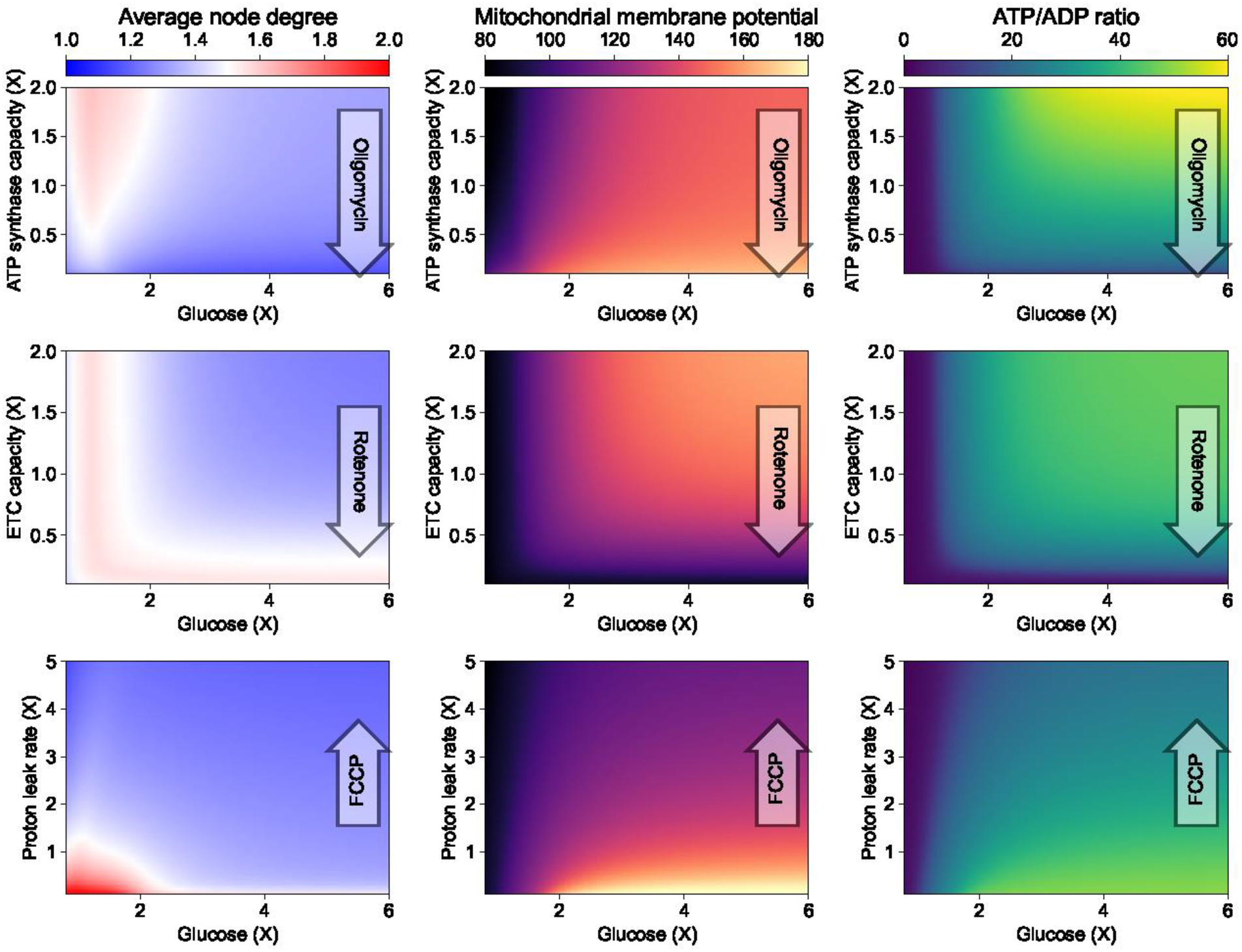
Steady-state values for a 2D parameter range of glucose concentrations and mitochondrial bioenergetics. The steady-state values of average node degree (first column), mitochondrial membrane potential (Δψ, second column) and ATP/ADP ratio (third column) are represented by the colors in the 2D contour plots. Relative glucose concentrations are in the X-axis; 5 mM is denoted as 1X. Relative activities of ATP synthase (first row), electron transport chain (ETC, second row), and proton leakage activity (third row) in the Y-axis are presented by comparing to the baseline model values. The translucent arrows indicate how the mitochondrial parameters change in response to the addition of oligomycin, rotenone, or FCCP.

**Table 1:**
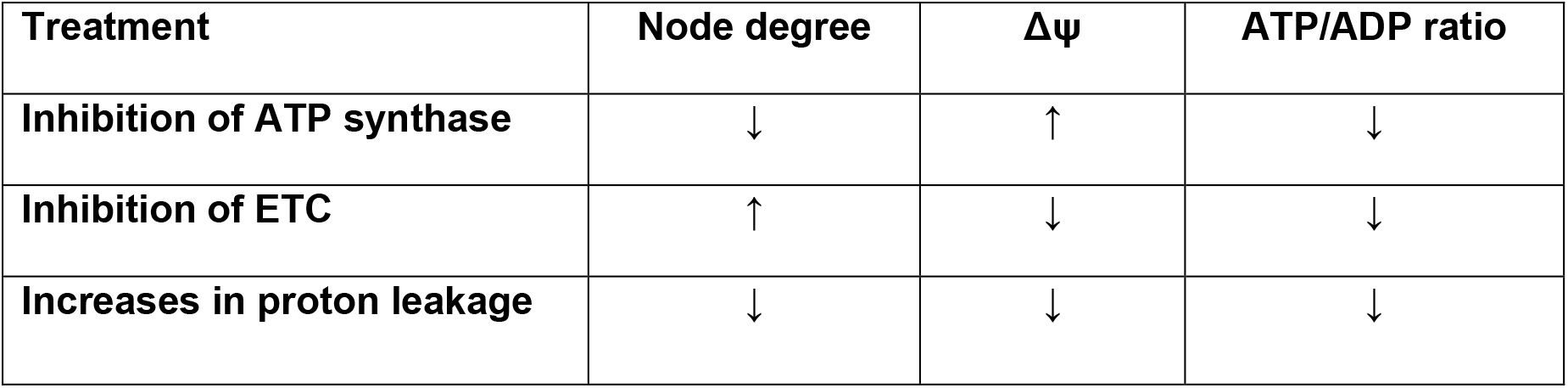
Summaries of the outcomes to bioenergetic change shown in Figure 3.

### Microscopic images of INS-1 cells presented a similar trend

To corroborate the findings from our *in silico* model, we obtained fluorescence microscopic images of INS-1 rat insulinoma cells under various metabolic conditions. First, we analyzed mitochondrial network morphologies under a range of glucose concentrations (0X, 1X, 3X, and 6X). We then performed a similar analysis with three different chemical agents: rotenone to block complex I in the electron transport chain (ETC), oligomycin to block ATP synthase, and trifluoromethoxy carbonyl cyanide phenylhydrazone (FCCP) to increase proton leakage across the IMM. The upper boxplots (Fig. 4 C) showed that the intensities of the voltage-dependent TMRM dye were the lowest at 0X, whereas the higher values were obtained under 1X, 3X, and 6X irradiation, revealing higher mitochondrial membrane potentials. A similar trend in mitochondrial membrane potential was also observed in PANC-1 cell line stained with a fluorescent mitochondrion-specific dye and analyzed by flow cytometry (data not shown). The analysis of mitochondrial morphology revealed that the networks showed a higher degree of fusion under the baseline glucose concentration of 1X (5 mM) and were more fragmented when the glucose concentrations were either higher (3X, 6X) or lower (0X).

**Fig. 4:**
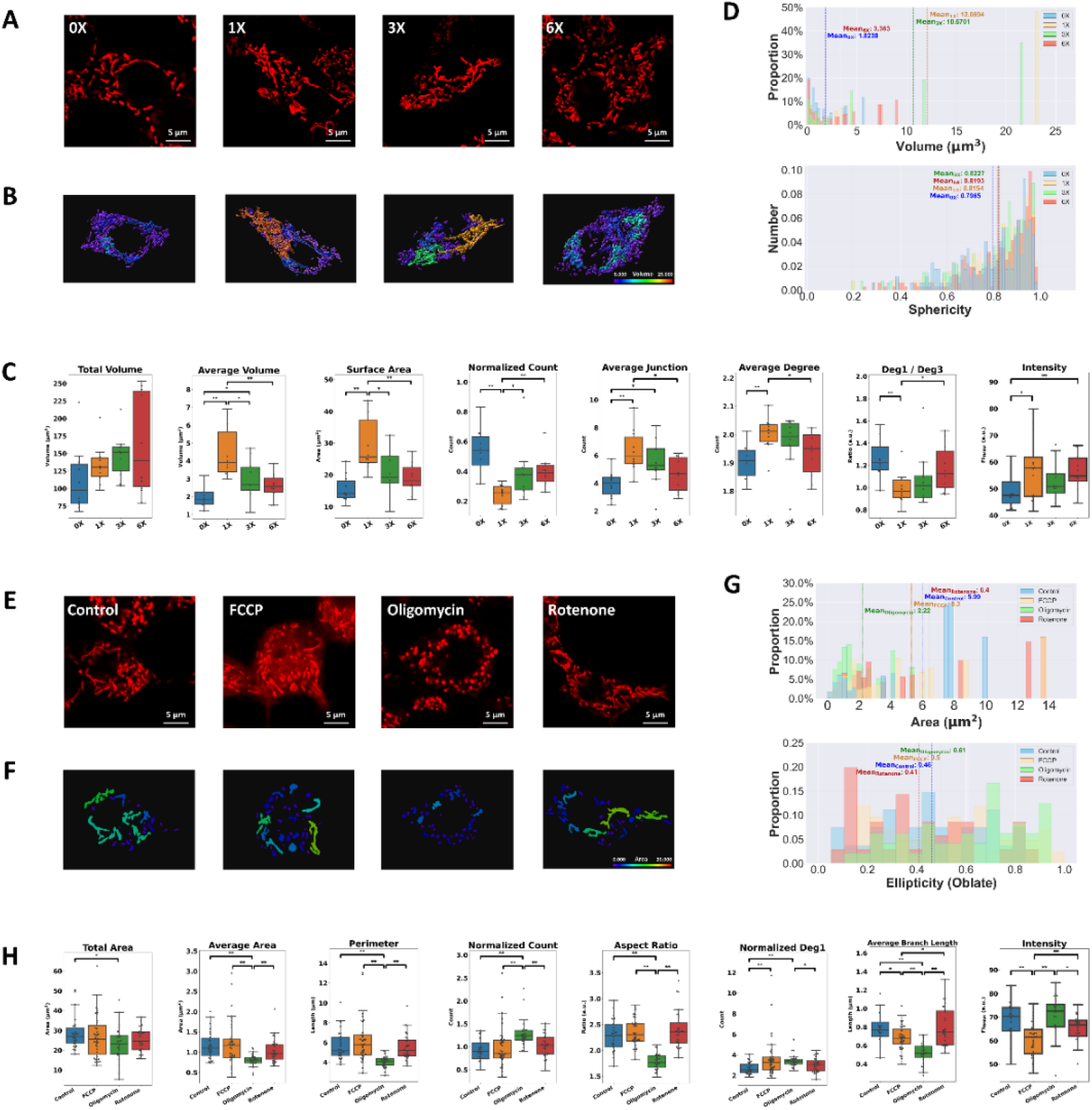
Mitochondria in INS-1 cells showed distinct morphologies under different cellular environmental conditions. (A) and (E): Representative fluorescent images of mitochondria in INS-1 cells labeled with TMRM under (A) four different glucose concentrations (with a baseline glucose concentration of 2 g/L, which is referred to as 1X) and (E) in the presence of three chemicals at a concentration of 10 μM. (B): Three-dimensional rendered surface images of the mitochondria in images (A). The images were created using Imaris software (Oxford Instruments). Separated mitochondria are labeled with different colors corresponding to their volumes. (F): Masked images of mitochondria in images (E). Binary images were first obtained using FIJI software and then mapped with different colors corresponding to their areas. (C) and (G): Histograms showing the distribution of different morphological indicators for separated mitochondrial components in an individual cell under several conditions. (C): Histograms of volume and sphericity. The data were acquired from cells in the images presented in (A). (G): Histograms of area and ellipticity. The data were acquired from cells in the images shown in (E). (D): Box plots of the 3D image analysis of mitochondria in INS-1 cells under different glucose concentrations. N = 10 cells for each concentration. (H): Box plots of the 2D image analysis of mitochondria in INS-1 cells under different chemical conditions. N = 26, 39, 28 and 31 for the control, FCCP, oligomycin and rotenone groups, respectively. All data for morphological indicators were obtained using the image analysis pipeline presented in the Materials and Methods section. Statistical significance was calculated using Levene’s test, one-way ANOVA, Student’s t-test, and Welch’s t-test (*: p<0.05, **: p<0.01, ***: p<0.001; n=10 cells for each).

The lower boxplots (Fig. 4 H) showed that mitochondria treated with oligomycin exhibited the highest membrane potential, whereas those treated with FCCP had the lowest potentials. In addition, the rotenone group had a slightly lower membrane potential than the control group, but this difference was not statistically significant. The results were consistent with the respective chemical mechanisms: oligomycin blocks ATP synthase, rotenone blocks the ETC, and FCCP increases proton leakage. With respect to the mitochondrial morphology, the oligomycin group showed the most fragmented mitochondrial networks and rounded mitochondria (Fig. 4 E). The FCCP group was less fragmented than the oligomycin group but more fragmented than the control group, although the lower pixel intensity might impact the quality of the analysis. The rotenone group showed an insignificant change in mitochondrial morphology compared with the control group.

### Prediction of the response to calcium oscillations

Calcium is one of the key players in insulin secretion by pancreatic cells. Mitochondria sense and shape the cytosolic calcium in response to glucose stimulation to modulate metabolism-secretion coupling. How mitochondria act as both recipients and generators of calcium signals would be an interesting factor to explore using our model. Two main types of cytosolic calcium oscillations have been observed in pancreatic β-cells: fast with the period ranges in seconds, and slow with periods of minutes[48]. In this study, we focus only on the slow oscillations in pancreatic β-cells. The cytosolic calcium levels in our *in silico* model are steady-state averages controlled solely by the cytosolic ATP/ADP ratio without any sophisticated electrophysiology and oscillations. Therefore, we used an independent periodic oscillator to complement the study to assess how the *in silico* model responds to oscillating cytosolic calcium levels. The shape and period of the calcium oscillator were made similar to one previous study [49] of mouse pancreatic β-cells under glucose stimulation.

The mitochondrial calcium levels oscillated with a larger amplitude and a slight delay compared with the cytosolic calcium levels. However, the levels of G3P, pyruvate, and NADH, the ATP/ADP ratio, and the mitochondrial membrane potential decreased as the level of cytosolic calcium increased and vice versa. The mitochondrial network tended to fuse as the cytosolic calcium level increased and slowly became more fragmented as the cytosolic calcium level decreased (Fig. 5).

**Fig. 5:**
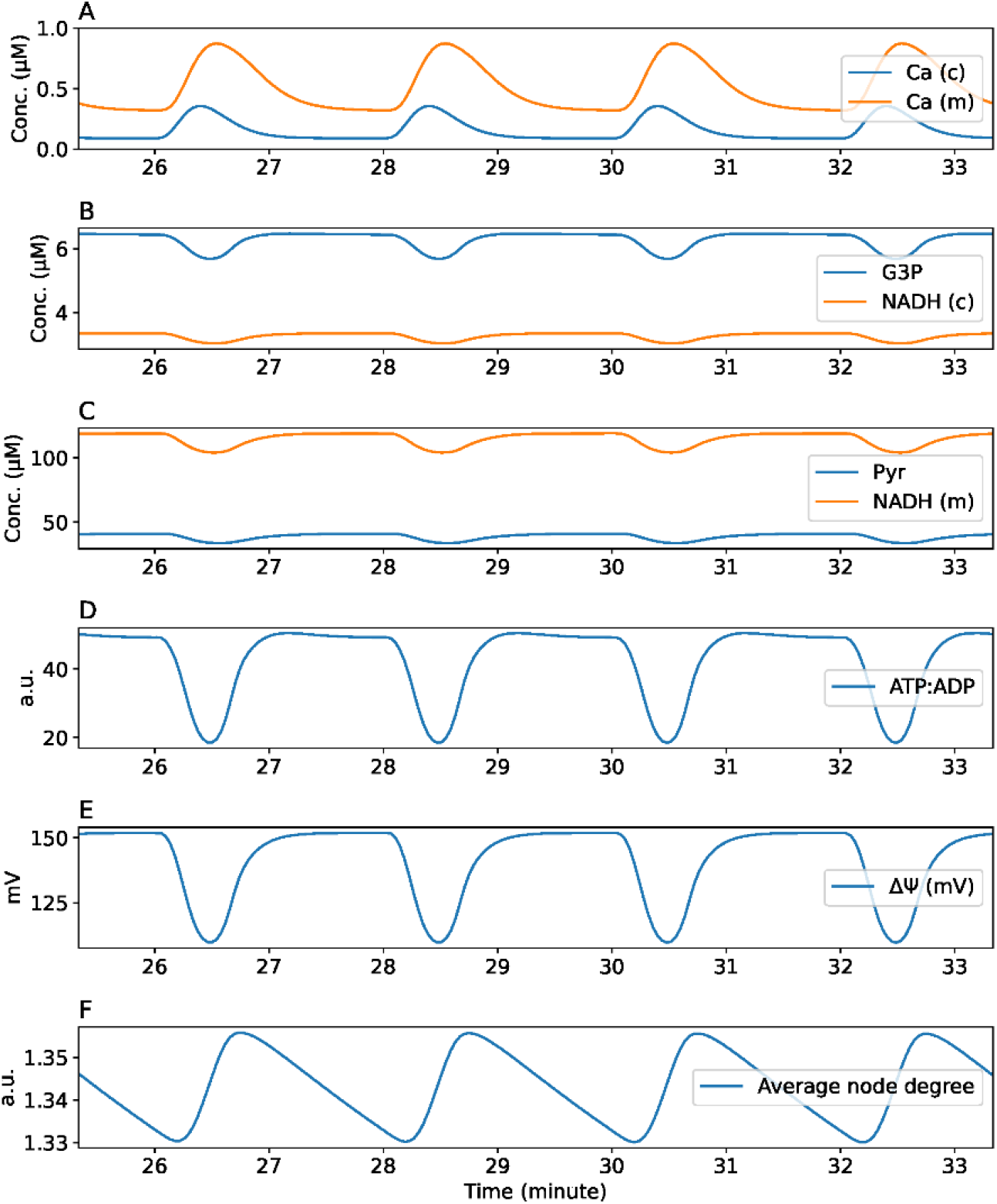
Response to cytosolic calcium oscillations. Levels of (A) cytosolic and mitochondrial calcium (in μM), (B) G3P and cytosolic NADH (in μM), (C) pyruvate and mitochondrial NADH (in μM), (D) ATP/ADP ratio, (E) mitochondrial membrane potential (in mV), and (F) average degree of nodes in the mitochondrial network were shown in last 8 minutes of cytosolic calcium oscillations.

### Predicting the blunted response of diabetic cells to glucose stimulation

To simulate the inhibition of mitochondrial metabolism in diabetic cells, we restricted the enzyme activities in the citric acid cycle (pyruvate dehydrogenase, PDH) and OXPHOS (ETC, ATP synthase)[18] and found that the proton leak rate was also increased [50] (Table 3). The diabetic condition was first tested by sequential challenges of glucose and chemicals (Fig. 6A) as the experimental protocol performed in the mouse islet [18]. The diabetic model had lower cytosolic NADH levels but markedly higher pyruvate levels at higher glucose concentrations than the baseline model (Fig. 6B (A), (B)). The diabetic model also showed an attenuated response to higher glucose levels in terms of ATP levels, cytosolic and mitochondrial calcium levels, and the mitochondrial membrane potential (Fig. 6B (E), (F), (G), (H)). Compared with the baseline model, the diabetic model presented higher mitochondrial NADH levels at lower glucose levels, but the baseline model showed higher mitochondrial NADH levels at higher glucose levels (Fig. 6B (D)). Across various glucose levels, the change in the mitochondrial network morphology was less prominent in the diabetic model than in the baseline model (Fig. 6B (I)).

**Fig. 6:**
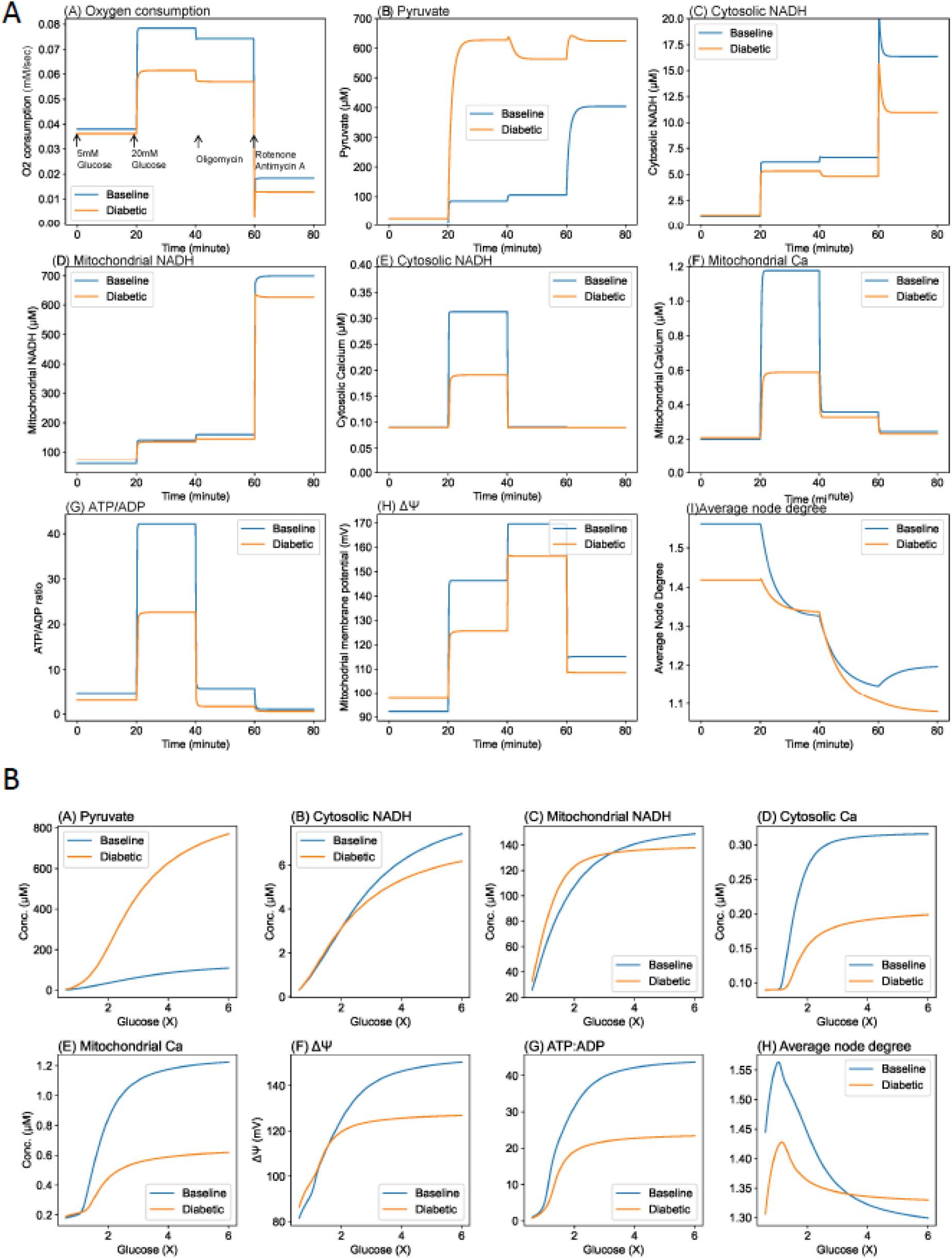
*In silico* ODE model’s response to glucose stimulation in baseline and diabetic models. A. Comparison of time-series responses to glucose and chemical stimulations in baseline and diabetic models. Oxygen consumption rate, metabolites, calcium, ATP/ADP ratio, mitochondrial membrane potential and average degree of nodes were compared followed by sequential additions of glucose and chemical reagents in baseline and diabetic models. Steady-states at the baseline glucose concentration of 5 mM were denoted at time t=0. At the time of 20 mins, glucose level was increased to 20 mM. At the time of 40 mins, ATP synthase activity was restricted to 5% of its original value to simulate oligomycin addition. At the time of 60 mins, ETC was restricted to 5% of its original value to simulate rotenone/antimycin addition. B. Glucose stimulation in baseline and diabetic cells. Baseline and diabetic cell steady-state variables of pyruvate, cytosolic and mitochondrial NADH and calcium, mitochondrial membrane potential, ATP/ADP ratio, average degree of nodes in the mitochondrial network were compared under different glucose concentrations. Relative glucose concentrations; 5 mM is presented as 1X.

## Discussion

Mitochondria are highly dynamic and motile organelles that undergo constant fusion and fission, as observed from live-cell fluorescent imaging. These fission/fusion processes are critical for mitochondrial quality control and function. An *in silico* model of mitochondrial bioenergetics and dynamics was proposed in this study, and this model reproduced several microscopic observations of the pancreatic cell mitochondrial network morphology under various glucose concentrations and chemical treatments. Increasing the glucose concentration, inhibiting ATP synthase, and increasing proton leakage resulted in a fragmented mitochondrial network. The *in silico* approach provides insights into the driving forces of mitochondrial dynamics: the fission rate was related to proton leakage, and the fusion rate was associated with ATP synthase.

### *In silico* model of bioenergetic influences on mitochondrial dynamics

Our *in silico* model was constructed based on two previous independent works that governed glucose-stimulated ATP synthesis [42] and fission/fusion processes of mitochondrial networks[46]. What bridges cellular bioenergetics and mitochondrial dynamics together is the choice of fission/fusion rates. For simplicity, we fixed the mitochondrial fission rate according to previous studies [16,44,47] and scaled the fusion rate to the ratio of the ATP synthesis (J_ANT_) rate and the proton leak rate (J_HL_). Choosing these two rates alone enabled us to simulate the mitochondrial network architectures in bioimages under different modes of energy supply and expenditure [5]. From the viewpoint of molecular biology, uncoupling (proton leak) activates calcineurin and then Drp1 [6,51] to trigger mitochondrial fission. In contrast, mitochondrial ATP synthesis could fuel Opa1 [6] through the actions of adenine nucleotide translocators (ANTs) and mitochondrial nucleoside diphosphate kinase (NDPK-D) [52,53], which enhances mitochondrial fusion.

From the viewpoint of bioenergetics, at the steady state, the energy provided by glycolysis and the citric acid cycle should balance the energy produced by ATP (via ATP synthesis) and dissipated as heat (e.g., via proton leak). In our model, this fact translates to the conservation law in the proton circuit: the protons pumped out of mitochondria by electron transport chain complexes (_JHRS_) should be equivalent to those returning to mitochondria to form the proton flux through ATP synthase (J_HF_) and leakage (J_HL_) through the IMM. In other words, at the steady-state,

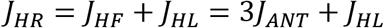

A higher proton leak rate (J_HL_) relative to the ATP synthesis rate (J_ANT_) results in more fragmented mitochondrial networks and a smaller average degree of nodes, and vice versa. High glucose concentration increases the mitochondrial membrane potential (Δψ) (Fig. 2), J_ANT_, and J_HL_. However, J_HL_ is exponentially dependent on Δψ and increases at a faster rate than the Hill equation relationship between J_ANT_ and Δψ (Fig. 7). Therefore, at higher glucose levels, the mitochondrial network was more fragmented with a smaller average degree of nodes. The mitochondrial network also became more fragmented at glucose levels lower than the baseline value of 1X (5 mM) because J_ANT_ decreased at a faster rate than J_HL_. At approximately the baseline glucose level (1X, 5 mM), the difference between fusion and fission rates was highest, and the most fused mitochondrial network was obtained. The inhibition of ATP synthase and increases in proton leakage resulted in a fragmented mitochondrial network. The model could predict observed mitochondrial fragmentation (Fig. 4) upon the addition of FCCP and oligomycin because the former enhances proton leakage (J_HL_) and the latter inhibits ATP synthase (J_ANT_) (Fig. 7). The addition of rotenone to block complex I, as simulated by reducing the ETC activity in the ODE model, hampered the response to glucose addition (Fig. 7) because the fission/fusion forces (J_HL_, J_ANT_) were both limited by the upstream reaction (J_HR_). In our study, rotenone caused insignificant changes to the mitochondrial network architectures in the INS-1 microscopic images (Fig. 4). However, the effects might depend on the dosage and cell type.[54,55]

**Fig. 7.**
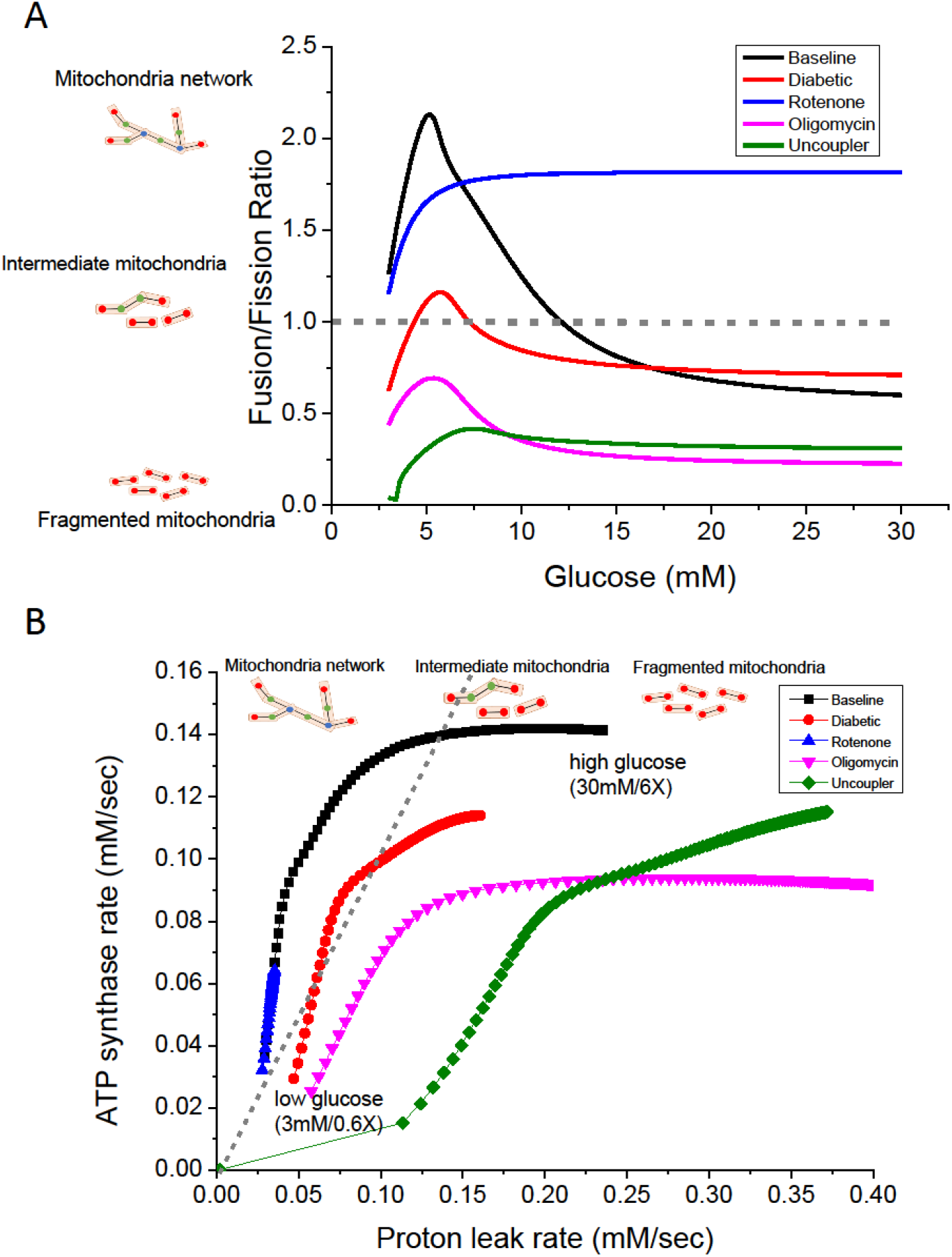
Effects of glucose stimulation under various conditions. A: Ratios of the fusion to fission rates under various conditions: default parameters (baseline, blue), diabetic parameters (diabetic, red), 90% ETC inhibition (rotenone, green), 90% ATP synthase inhibition (oligomycin, cyan), and five times the proton leak rate (Uncoupler, black). B: Steady-state proton leak rate (fission force) and ATP synthesis rate (fusion force) under various conditions (color codes are identical to A). Each dot represents the glucose concentration starting from 3 millimolar (the lower left ones) to 30 millimolar (the upper right ones) with an increment of 1 millimolar.

Overall, perturbations to mitochondrial energetics, such as increasing the proton leak rate, inhibiting ATP synthase, and combined parameter alterations in diabetic cells, made the mitochondria more fragmented. In contrast, inhibiting the ETC yielded a more fused mitochondrial network at high glucose concentrations because limiting proton pumping restricted JHL more than JANT. Compared with default parameters, perturbations to mitochondrial energetics parameters also reduced the dynamic range of fission/fusion rates across various glucose concentrations, which implied that the mitochondrial network structure is less responsive to metabolic changes. Our results also confirmed that the inhibition of mitochondrial fusion does not necessarily require a loss of the mitochondrial membrane potential but depends on the proton motive force.

### Extending the model to diabetic beta cells

Mitochondrial dysfunction, including changes in respiratory chain activity, is related to dysfunctional glucose-stimulated insulin secretion (GSIS) in models of type 2 diabetes. Energy changes, such as reduced ATP levels, ATP/ADP ratios and mitochondrial membrane potential, were observed in diabetic beta-cells [56], and these cells showed increased of mitochondrial complex I, ATP synthase, UCP-2, and reactive oxygen species [57]. While mitochondrial dynamics maintain the metabolically efficient mitochondrial population, an imbalance between mitochondrial fusion and fission also contributes to the beta-cell dysfunction in the progression of diabetes.

In the diabetic cell simulation, we restricted the activities of several metabolic checkpoints, including pyruvate dehydrogenase (PDH), which controls the citric acid cycle (CAC), the electron transport chain (ETC), and ATP synthase, to reflect inhibited mitochondrial metabolism in diabetic cells.[18] In our simulations, the mitochondrial membrane potential and NADH levels were higher in the diabetic model than in the baseline model at a resting glucose concentration of 5 mM (Fig. 6B (C) and (F)). However, the reverse was observed with increases in the glucose concentration. This phenomenon was also observed by Haythorne et al.[18]: the restriction of mitochondrial metabolism blunted the responses to increasing glucose levels across various variables starting from the CAC, including mitochondrial calcium, NADH, and membrane potential. The limited ATP synthesis obtained from decreased ATP synthase activity and a lower mitochondrial membrane potential hampered the increases in the cytosolic ATP/ADP ratio and the cytosolic calcium levels (Fig. 6A, Fig. 6B (D), (G)), reflecting less effective insulin secretion in response to glucose stimulation. While we increased the basal proton leak rate in the diabetic model in our simulation (Table 3), the contribution of proton leak in total oxygen consumption rate (OCR) did not show much differences between healthy and diabetic (Fig 6A (A)) as observed in the previous study of mouse islets by Haythorne et al. [18]. The sensitivity to glucose stimulation in the ODE model mainly relied on the proton leak rate [42]. A larger basal proton leakage rate would shift the curves of calcium and ATP response rightwards (Fig. 6B), making the cells less responsive to glucose stimulation and more consistent with diabetic conditions, including persistent high glucose and lipid levels. Additionally, proton leakage across the inner mitochondrial membrane is mediated via uncoupler protein 2 (UCP2), which is stimulated by high nutrient levels and ROS production, and disrupts the mitochondrial membrane potential, ATP synthesis, and insulin secretion.[50]

With respect to the mitochondrial network architecture, diabetic cells showed a more fragmented configuration than baseline cells in the model at a resting glucose concentration of 5 mM, as implied by a smaller average degree of mitochondrial nodes (Fig. 6), which was consistent with previous microscopic observations.[20,23,58] The range of the average degree of mitochondrial nodes across various glucose concentrations was less prominent in the diabetic model than in the baseline model, which might reflect that diabetic beta cells exhibit less metabolic plasticity than healthy beta cells (Fig. 6B (H), Fig. 7).[5,51] Aging also shows a similar metabolic syndrome characterized by impairments in glucose homeostasis, mitochondrial metabolism and insulin secretion from pancreatic β-cells.[48] Our model might be able to explain the mitochondrial morphology, which is prone to become swollen and fragmented with age. [59,60]

### Analysis of fission/fusion events

To obtain a more detailed understanding of the relationship between fusion/fission rates and environmental conditions, we adapted the graph-theory-based network model[46] to realize mitochondrial fission/fusion processes in individual mitochondria (Fig. 8). The network model consists of two types of fusion and fission events: tip-to-tip (C1) and tip-to-side (C2) fusion to grow and branch out the mitochondrial network. By observing C1 and C2 extracted and fitted to microscopic images from the mitochondrial network model, we could estimate the propensity of different types of mitochondrial fusion and fission events under different conditions. In the case of low to high glucose conditions, mitochondria under 1X and 3X concentrations harbored both higher C1 and C2, which was consistent with the network-like morphologies observed from the image analysis. The lower values of C1 and C2 at 0X also corroborate the fragmented morphology of mitochondria. Additionally, the mitochondria at 6X harbored a high value of C1, but the low C2 was similar to that at 0X, which provided a clue to the hypothesis that the fragmentation of mitochondria observed at high glucose concentrations has a closer relationship to tip-to-side fission than tip-to-tip fission. In contrast, in toxicity fitting, significantly fragmented mitochondria induced by oligomycin presented a lower C1 value, and higher C1 values were found in the control group and in the group treated with rotenone, showing consistency between the image analysis results and the parameter fitting obtained from network modeling. The C2 value varies over a wide range among samples even under the same conditions, which may result from the few degree-3 nodes in all toxic conditions, which makes C2 unstable during network model stimulation and GA fitting. By re-visualizing the mitochondrial dynamics in the agent-based model simulation, “Average Degree” were tracked along with the iterations under different conditions, where fragmented networks such as those under oligomycin and 0X conditions owned smaller values during simulations. Harboring the largest C1 and C2, a simulated model for 1X showed the most complicated network. In contrast, a simulated network for 0X was significantly fragmented with more segmented short branches, which was consistent with the results of image analysis and GA fitting.

**Fig. 8:**
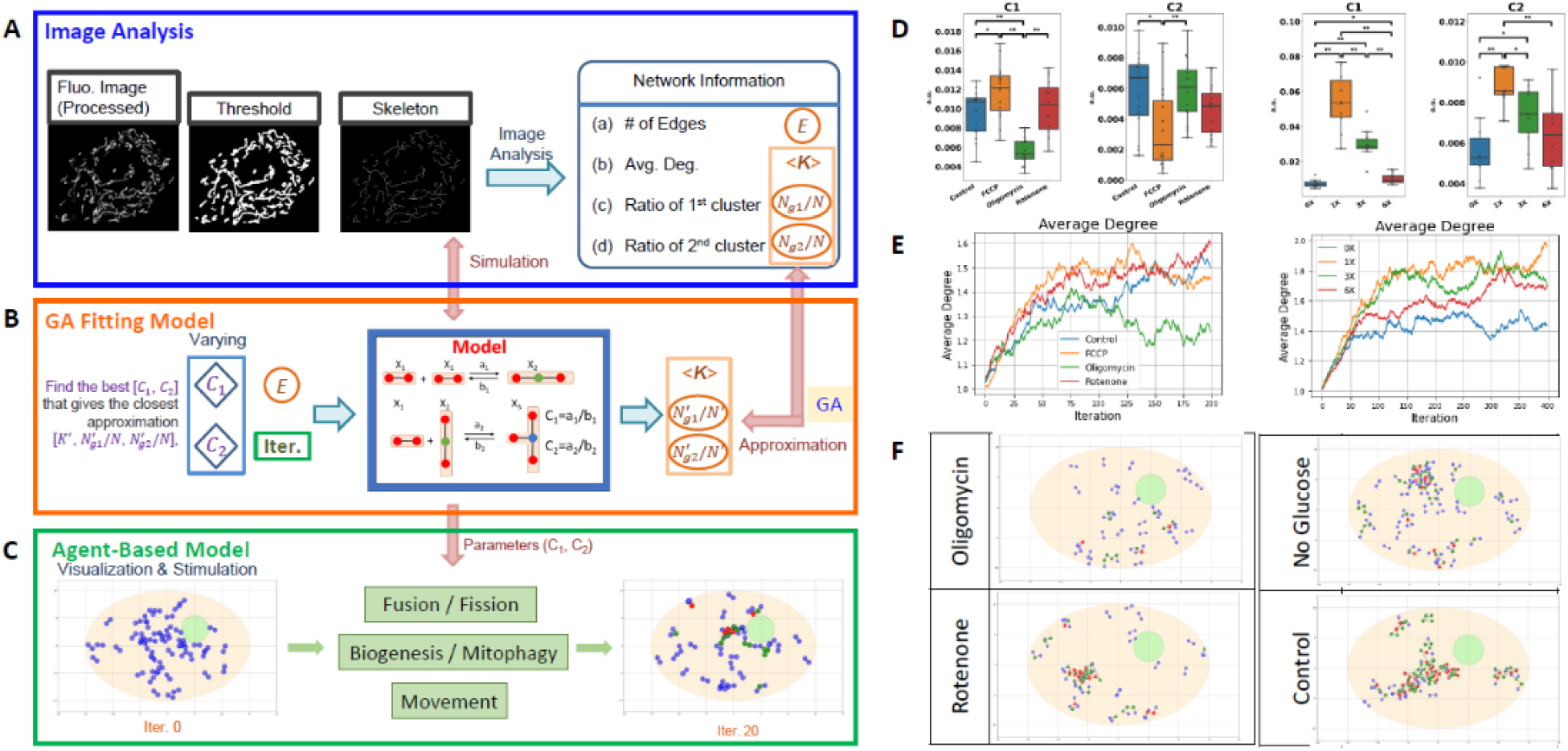
Mitochondrial network model stimulation under various metabolic conditions. A: Workflow of mitochondrial network model simulation and fitting fission/fusion rates. The agent-based mitochondrial network model simulating two types of fission and fusion behaviors using nodes, edges, and degrees are used to represent and describe the mitochondrial network. Edges are considered as the basic units which represent small segments of mitochondria, and nodes with different degrees are regarded as characteristic measurements of how fragmented the mitochondrial network is. B. Fitting fission/fusion rates of mitochondrial under different glucose concentrations and chemical treatment. Network parameters <k> (average degree), Ng1/N (number of nodes or edges of the largest cluster / total nodes or edges), Ng2/N (number of nodes or edges of the second largest cluster / total nodes or edges) were extracted from fluorescent images of INS-1 and used as features for GA (genetic algorithm) fittings. By minimizing the distance between the distribution of network parameters (D(<k>, Ng1/N, Ng2/N)) extracted from fluorescent image analysis and the network model, the optimized C1 (ratio of the rate constants of tip-to-tip fusion to tip-to-tip fission) and C2 (ratio of the rate constants of tip-to-side fusion to tip-to-side fission) were obtained by random search of GA. Kernel Density Estimation (KDE) was used to estimate the probability density function of the network parameters from confocal microscopy images of mitochondria, and Kullback-Leibler Divergence (KLD) was used to minimize the difference between two distributions calculated from KDE. C. Agent-based model for visualization of the simulated mitochondrial networks with C1 and C2 obtained from GA. D. Ten and fifteen repeats of fitting for glucose and toxicity 2D data respectively. E. Tracking indicator “Average Degree” of simulated networks along the iterations. F. Snapshots of realization of mitochondrial network in the agent-based model.

### Future works

All models are abstractions to their real-world counterparts, and our ODE model is no exception. Our ODE model emphasizes the influence of mitochondrial bioenergetics on mitochondrial morphology. We made some assumptions and simplifications in the model to make it easy to constrain the parameters and analyze the general behavior of the model.

In our model, glucose served as the sole energy source. Glycolysis and OXPHOS were the main metabolic pathways driving GSIS. Several related metabolic pathways were not explicitly covered in the model, such as the pentose phosphate pathway (PPP)[3,18], reactive oxygen species (ROS) generation and signaling[55,61], amino acid metabolism[13,18,22,62–64], lipid homeostasis[22], and the mitochondrial GTP (mtGTP) cycle[65].

The cytosolic calcium levels in our model were steady-state averages as a function of the intracellular ATP/ADP ratio. Because the focus of this study is the steady-state analysis of metabolic (ATP, mitochondrial membrane potential) implications for mitochondrial fission/fusion dynamics, a steady-state average cytosolic calcium level would ease the study of model behaviors. To avoid introducing too much complexity into the model, the section representing calcium oscillations used an independent periodic function (Fig. 5 and Materials and Methods) rather than including full descriptions of plasma membrane electrophysiology and calcium signaling [41,66,67]. This simple modeling could still reproduce the curves of cytosolic calcium and ATP levels in the previous beta-cell study.[49] The average degree of mitochondrial nodes fluctuated due to varying ATP synthesis rates and mitochondrial membrane potentials, albeit at a smaller amplitude because the time scale of fission/fusion dynamics was substantially longer (10 minutes) than that of calcium oscillations (2 minutes).

In addition to ATP synthesis and proton leakage flux, several bioenergetic mechanisms are also involved in mitochondrial fusion/fission cycles.[6] For instance, the fusion GTPase Opa1 degrades in depolarized mitochondria to prevent damaged mitochondria from merging with the mitochondrial network.[68] Another component not included in this model was AMP-activated kinase (AMPK). During starvation, an increased AMP/ATP ratio activates AMPK and triggers downstream signal transduction pathways to promote mitochondrial fission, mitophagy, and biogenesis [69–71]. However, the mitochondrial mass was considered constant in our model without mitophagy or biogenesis processes. Additionally, the simulations focused on glucose levels higher than the baseline value of 5 mM; therefore, AMPK activation via starvation played a lesser role in our simulations.

The mitochondria were assumed to be well mixed in our model with adequate fusion events[46,68]. Each mitochondrial node had the same metabolic activities. Therefore, the mitochondrial dynamics part was simplified to only two ODEs (populations of degree-2 and degree-3 nodes, respectively). The average degree of nodes could represent the general trend of the mitochondrial network, regardless of whether it is fused or fragmented. However, an agent-based approach[16,44] is needed to trace individual mitochondria to monitor the health status and mitochondrial cluster formation. Each mitochondrial particle cost three state variables (NADH, calcium, and the mitochondrial membrane potential) in the ODE model. The added complexity is beyond the scope of this study and could be included in future work.

Mitochondrial network organization and bioenergetic functions have bidirectional relationships[54]. Mitochondrial fission/fusion affects bioenergetic efficiency and energy expenditure, whereas the mitochondrial morphology changes according to the cellular energy state. The metabolic signaling pathways AMPK, insulin/IGF, and mTOR regulate mitochondrial dynamics for structural and functional adaptation. The bidirectional relationship and the detailed regulatory mechanisms still need further elucidation. Nonetheless, our simplified ODE model captured the changes in the mitochondrial network morphology under different metabolic conditions and could serve as the basis for future works.

In conclusion, we devised a simple ODE-based mathematical model to bridge cellular bioenergetics and mitochondrial dynamics, and this model corroborated the behaviors based on fluorescence microscopy findings from INS-1 cells. The model also demonstrated that mitochondrial dynamics were regulated by ATP synthesis and proton leakage under various metabolic conditions. The mitochondrial networks were reconstructed in the agent-based simulations that incorporate the fission/fusion rates from the model and image analysis. The combination of biophysical modeling and network analysis on experimental data can provide insights into the fundamental principles underlying these complex regulations of organelle dynamics.

## Material and Methods

### *In silico* ODE model

Our computer simulation model (Fig. 1) was built upon ten coupled ordinary differential equations (ODEs) that integrate the bioenergetic and mitochondrial dynamics parts of the model. The former was extended from the glucose-sensing beta-cell model by Fridlyand et al. [42], and the latter was derived from the graph theory-based mitochondrial network model by Sukhorukov et al. [20,46,72]. The bioenergetic part described the biochemical reactions of glycolysis, the citric acid cycle (CAC), oxidative phosphorylation (OXPHOS), and calcium dynamics. First, glucose is metabolized to pyruvate via glycolysis. Pyruvate is then consumed in the mitochondria in the citric acid cycle (CAC) to generate the reducing equivalents NADH and FADH2, which fuel electron transport chain (ETC) complexes to pump hydrogen ions out of mitochondria and thus create a voltage and pH difference across the inner mitochondrial membrane (IMM), which is known as the proton motive force (PMF). The PMF is then tapped by F1-Fo ATPase (ATP synthase) to synthesize ATP. The increase in the ATP/ADP ratio triggers calcium influx into the cytosol and then into the mitochondria through the mitochondrial calcium uniporter (MCU), which enhances the CAC and OXPHOS reaction rates to generate even more ATP. The influx of calcium into the cytosol also triggers the exocytosis of insulin-containing vesicles. The entire biological process is called glucose-stimulated insulin secretion (GSIS).

Compared with the original work performed by Fridlyand et al. [42], we adjusted the mathematical expressions of adenine nucleotide translocator (ANT), mitochondrial sodium-calcium exchanger (NCLX) [31,73], and added adenylate kinase (AdK) equilibria[74] for the cytosolic ATP-ADP-AMP pool. All the changes were described in the Supplementary Material. Glucose-driven aerobic glycolysis was the sole energy source; metabolic pathways for amino acids and lipids were not included.[13,75] The mitochondrial dynamics section describes the fission-fusion cycles of mitochondrial nodes and segments. The movement and fission/fusion process of each mitochondrion were not tracked individually, similar to the process in agent-based modeling. We assumed that the mitochondrial mass was conserved and uniform in this model. The mitochondrial networks were represented by edges (mitochondrial mass) and nodes of degree 1 (end nodes), degree 2 (line segment nodes), and degree 3 (branching nodes). The fission/fusion cycle could be shown by the merging/splitting of the nodes (Fig. 1). The mitochondrial fission rate was fixed once every 10 minutes[16], and the fusion rates were scaled against the ATP synthase rate and the proton leakage rate[5]. At a resting glucose concentration of 5 mM (denoted as 1X), our *in silico* ODE model reached a steady state consistent with that of the original GSIS model (Table 2). This ensures that the alterations in our ODE model did not induce marked deviations from the original behavior.

**Table 2:** Steady-state values of the computational model with a baseline glucose concentration of 5 mM. The complete mathematical descriptions for the *in silico* model are described in the Supplementary Material.

| State variables | Fridlyand's model | Our model |
| --- | --- | --- |
| G3P | 2.79 $\mu\text{M}$ | 2.90 $\mu\text{M}$ |
| Pyruvate | 8.62 $\mu\text{M}$ | 8.75 $\mu\text{M}$ |
| Cytosolic NADH | 0.97 $\mu\text{M}$ | 0.982 $\mu\text{M}$ |
| Mitochondrial NADH | 60 $\mu\text{M}$ | 57.3 $\mu\text{M}$ |
| ATP/ADP ratio | 4.6 | 4.75 |
| Mitochondrial calcium | 0.242 $\mu\text{M}$ | 0.202 $\mu\text{M}$ |
| Mitochondrial membrane potential ($\Delta\Psi$) | 94.7 mV | 92.2 mV |
| Degree-1 node population | N/A | 0.338 |
| Degree-2 node population | N/A | 0.303 |
| Degree-3 node population | N/A | 0.01815 |

#### Computational environments

The *in silico* model was written in Julia [76] and was run on a workstation with two 8-core Xeon E5–2620 v4 CPUs or on machines with multiple Intel Xeon Processors in TWCC cloud service (https://www.twcc.ai) for high-speed parallel computing. The ordinary differential equations (ODEs) were solved by the DifferentialEquations.jl [77] package and were visualized by the PyPlot.jl package, a Julia wrapper to the Python matplotlib[78] visualization package.

#### Steady-state values under a range of glucose concentrations

First, steady-state values were obtained with a baseline glucose concentration of 5 mM. The model was then simulated under a range of glucose concentrations from 3 mM to 25 mM until the model reached a steady state. We measured the influence of glycolytic flux on the steady-state levels of G3P, pyruvate, calcium, ATP, the mitochondrial membrane potential, the mitochondrial fission-fusion rates, and the state of the mitochondrial network.

#### Interactions of mitochondrial energetics and dynamics

The interactions of mitochondrial energetics and dynamics were demonstrated by varying the glucose concentrations and one mitochondrial parameter, namely, ATP synthase activity, ETC complex activity, or proton leakage rate.

#### Response to calcium oscillations

The cytosolic calcium in the model represented the steady-state levels controlled by the cellular ATP/ADP ratio. However, in this section, the cytosolic calcium concentration was independent of the ATP/ADP ratio and described by a time-dependent oscillator:

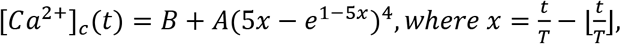

where A is the amplitude of the cytosolic calcium concentrations, B is the resting level of calcium, and T is the period of the calcium oscillator.

#### Mitochondrial dynamics in diabetic cells

We then compared how diabetic beta cells react to increases in glucose in terms of bioenergetics and mitochondrial dynamics. The capacities of pyruvate dehydrogenase (PDH), proton leakage, electron transport chain, and ATP synthase were altered in the simulation model to account for metabolic changes in diabetic beta cells. The steady-state values obtained from both settings were collected across a range of glucose levels from 3 mM to 30 mM.

**Table 3.**
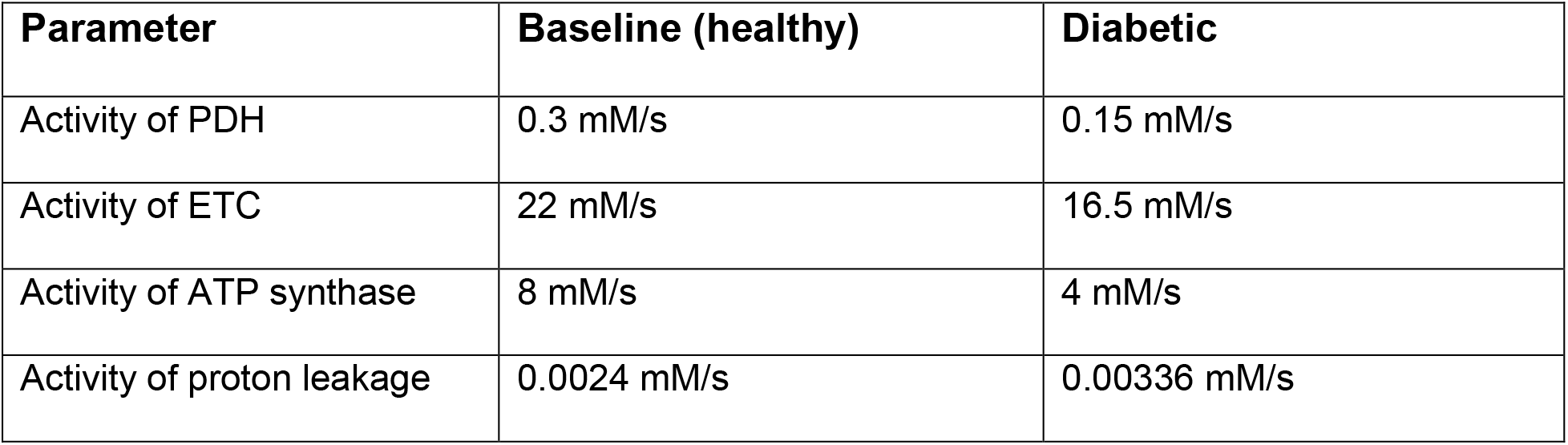
Parameters adjusted for diabetic beta cells compared with the baseline.

| Parameter | Baseline (healthy) | Diabetic |
| --- | --- | --- |
| Activity of PDH | 0.3 mM/s | 0.15 mM/s |
| Activity of ETC | 22 mM/s | 16.5 mM/s |
| Activity of ATP synthase | 8 mM/s | 4 mM/s |
| Activity of proton leakage | 0.0024 mM/s | 0.00336 mM/s |

### Microscopic Image analysis pipeline

#### INS-1 cell culture

INS-1 cells (INS-1 832/3 rat insulinoma cell line) were cultured in RPMI-1640 (Sigma Cat. No. R0883) supplemented with 2 mM L-glutamine, 1 mM sodium pyruvate, 10 mM HEPES (Cat. No. TMS-003-C), and 0.05 mM β-mercaptoethanol + 10% FBS at 37°C in the presence of 5% CO2 for 1 to 3 days before imaging.

For the glucose experiments, DMEM (no glucose, Gibco A1443001) was used as the culture medium after the previous medium was washed out with PBS. The culture glucose concentrations were 0 g/L (≈ 0 mM), 2 g/L (≈ 11.1 mM), 6 g/L (≈ 33.3 mM), and 12 g/L (≈ 88.8 mM), which are referred to as 0X, 1X, 3X, and 6X. Microscopic images were taken 30 minutes after the addition of DMEM and glucose.

For the toxicity experiments, the culturing media were the same as those used in the glucose experiment. Carbonyl cyanide-p-trifluoromethoxyphenylhydrazone (FCCP), oligomycin, and rotenone were added at a concentration of 10 μM. Microscopic images were taken immediately after the addition of FCCP, and in other cases, images were taken 30 minutes after addition of the chemicals.

Mitochondria were labeled with 100 nM tetramethylrhodamine, methyl ester (TMRM) and 100 nM nonyl acridine orange (NAO) for 15 minutes before imaging. A ZEISS LSM800 with Airyscan and a 1.40-NA 63x objective was used for cell imaging.

#### Image preprocessing and thresholding

The image analysis pipeline to obtain the mitochondrial network morphology information was based on the study performed by Chaudhry et al. [26] using the Fiji distribution in ImageJ2 software (Fig. 9). The inputs were TMRM or NAO fluorescence microscopic images of INS-1 cells. The pipeline comprised several preprocessing and analysis procedures. In the standard pipeline, input images were preprocessed by algorithms such as “Subtract Background” and “Enhance Local Contract (CLAHE)” to denoise and enhance the contrast. After preprocessing, the images were binarized and skeletonized to analyze the mitochondrial network morphology (Table 4).

**Fig. 9.**
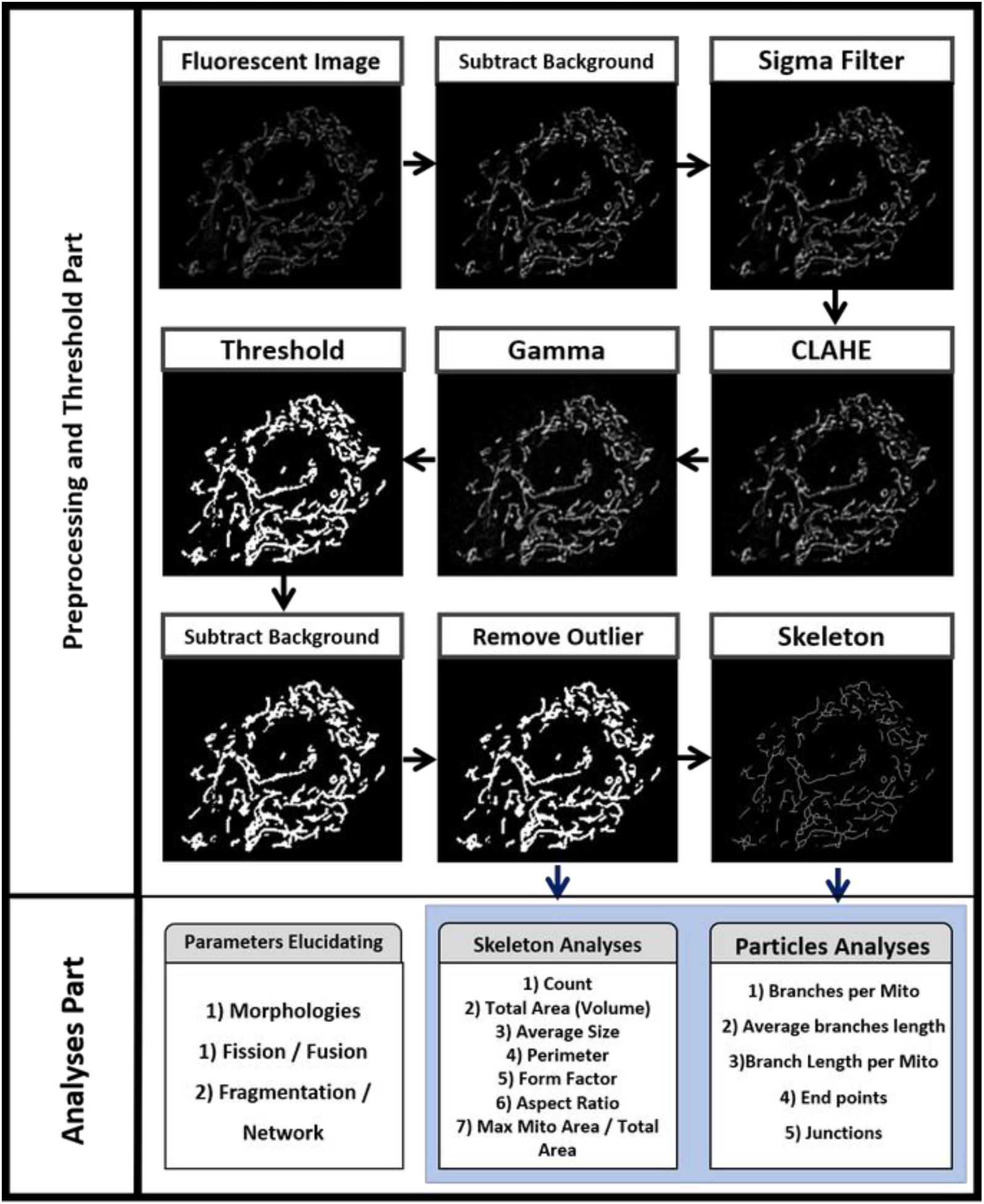
Workflow of the image processing pipeline for mitochondrial fluorescence images. Our pipeline is composed of the “Preprocessing and Threshold Part” and the “Analysis Part”. In the “Preprocessing and Threshold Part”, we applied several commands and algorithms from FIJI to obtain binary images and skeleton images, which would serve as the input images for the “Analysis Part”. Mitochondrial features were analyzed and calculated in the “Analysis Part” and were used as quantitative evidence for elucidating the morphologies and fission/fusion propensity of the mitochondria under certain conditions. For “optional adjustment” in the “Preprocessing and Threshold Part”, “add noise” and “brightness and contrast” could be applied according to the quality of the images and the thresholding performance.

**Table 4.**
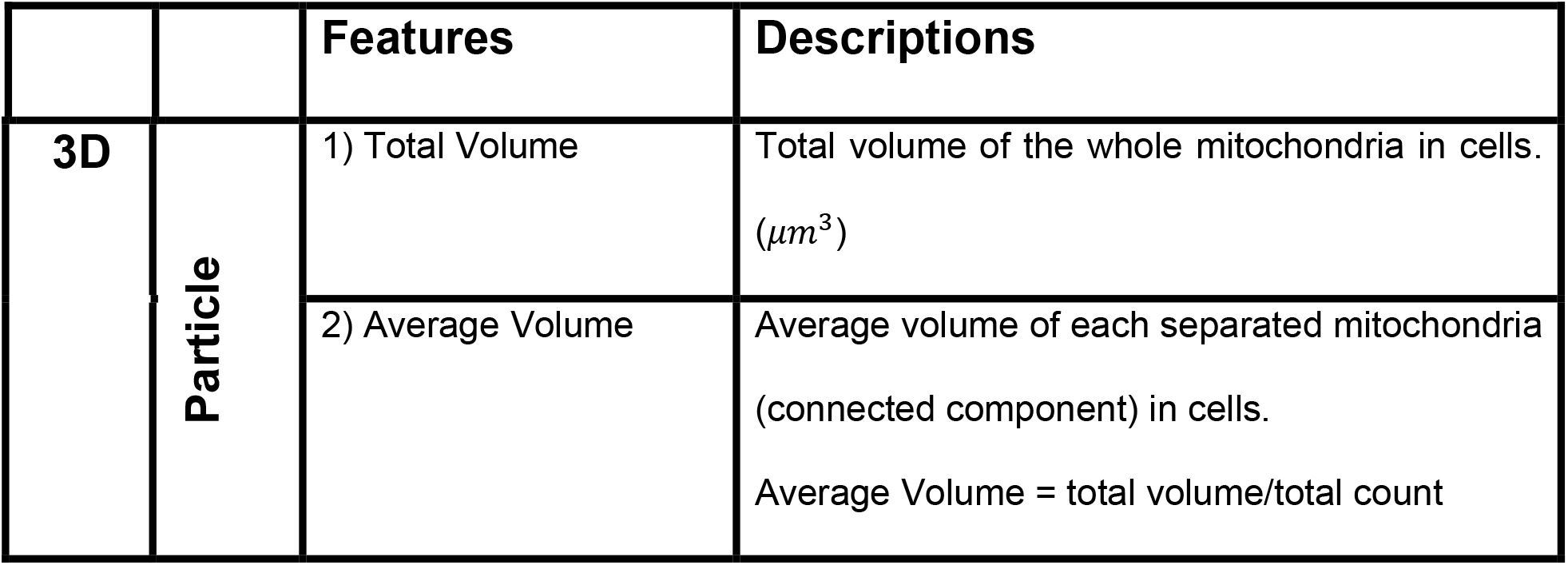
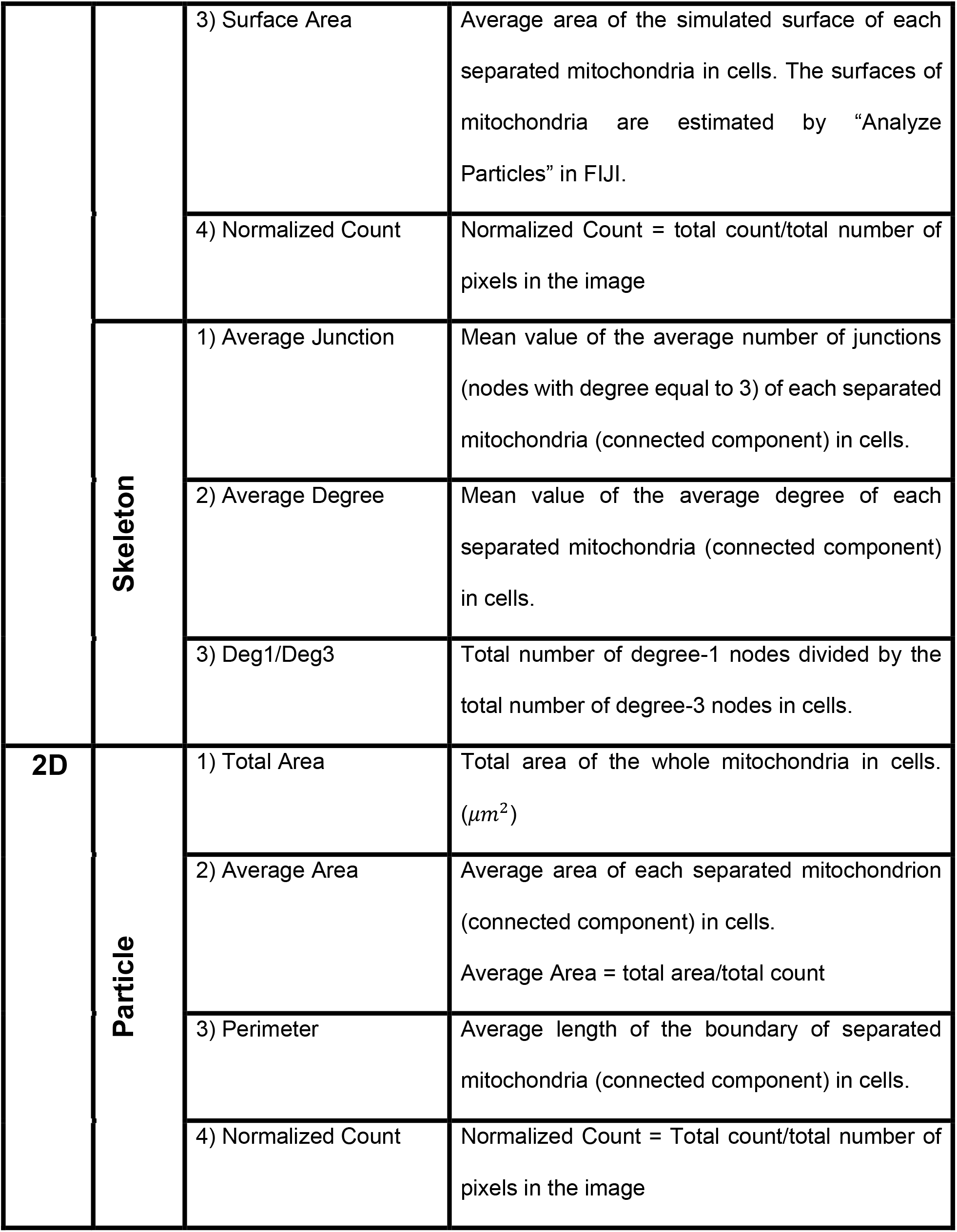
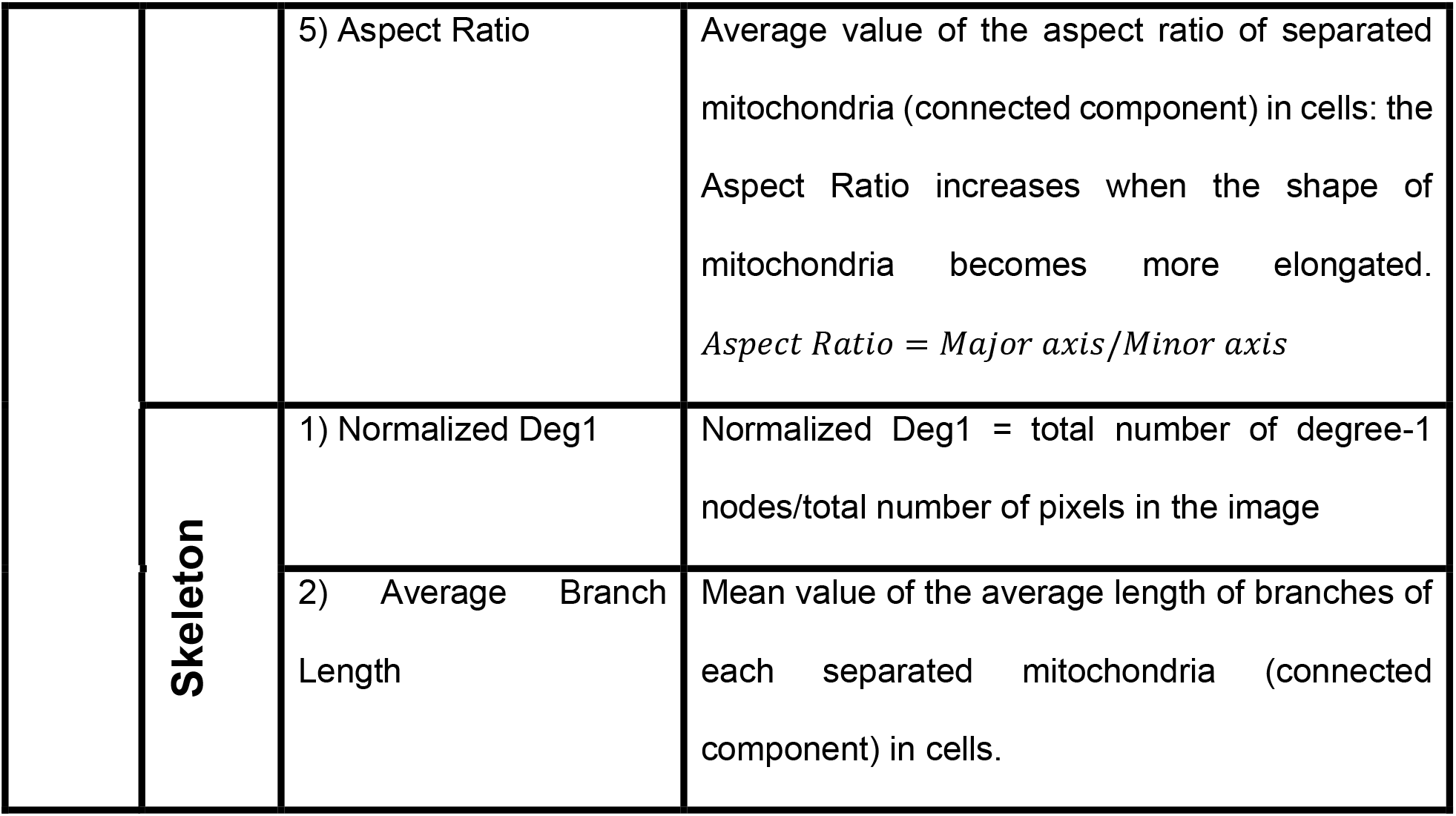
Description of mitochondrial features. The mitochondrial features consisted of two types: “particle analysis”, features derived from thresholded binary images, and “skeleton analysis”, features derived from skeletonized images.

#### Membrane potential analysis

The TMRM fluorescent intensity is an indicator of the mitochondrial membrane potential. We quantified the average pixel intensities of each TMRM image using ImageJ to evaluate the changes in the membrane potential under different glucose concentrations and chemical environments. The mean value of the “Auto Threshold” values (obtained from FIJI) of the control group is set as the threshold. We only considered pixels with intensities larger than the threshold to avoid interference by background noise.

#### Statistical analysis

One-way ANOVA was performed to analyze the data from the glucose doseresponse experiments. Welch’s t-test was applied to analyze the significant differences using the “ttest_ind” function from the “scipy.stats” package of Python. All data presented as boxplots were plotted using the “boxplot” function provided in the “matplotlib.pyplot” package of Python. The lower and upper bonds of the box were the first and third quartiles (Q1 and Q3) of the data, and the median (Q2) was the line inside the box. The lower whisker extended to the smallest data point being greater than Q1–1.5(Q3-Q1), and the higher whisker extended to the largest datapoint being smaller than Q3+-1.5(Q3-Q1). Statistical significance thresholds were set as *p<0.05, **p<0.01, and ***p<0.001.

## Acknowledgment

We thank to National Center for High-performance Computing (NCHC) in Taiwan for providing computational and storage resources.

## Funding

The work was supported by grants from the Ministry of Science and Technology in Taiwan (MOST-108-2636-B-002-001 and MOST-109-2636-B-002-001 grants to AW), and the grants from the Research Grants Council of the Hong Kong Special Administrative Region, China (Project #: CUHK 14201317 and C5011-19GF to YH) and the VC Discretionary Fund, the Chinese University of Hong Kong (Project #: 8601014 to YH).

## Data and code availability

The model code is available at https://github.com/NTUMitoLab/MitochondrialDynamics.

## Notes

### Competing Interest Statement

The authors have declared no competing interest.

