## Supplement for "Metabolic Regulation of Mitochondrial Morphologies in Pancreatic Beta Cells: Bioenergetics-Mitochondrial Dynamics Coupling"

### Supporting information

#### *In silico* ODE model descriptions

##### General parameters

| Parameter | Value | Description | Ref. |
|---|---|---|---|
| $V_i$ | 0.53 | Relative cytoplasmic volume | (1) |
| $V_m$ | 0.06 | Relative mitochondrial volume | (1) |
| $V_{mtx}$ | 0.0144 | Relative mitochondrial matrix volume (Adjustable) | (1) |
| $C_{mito}$ | 1.812 mM/V | Mitochondrial membrane capacitance | (1) |
| $F$ | 96484.6 C/mol | Faraday's constant | Physical constant |
| $\delta_{Ca}$ | 0.0003 | Fraction of free calcium in mitochondria | (1) |
| $[Na^+]_c$ | 10 mM | Cytoplasmic Na concentration | (1) |
| $[Na^+]_m$ | 5 mM | Mitochondrial Na concentration | (1) |
| $T_v$ | 26.73 mV | Thermal voltage (RT/F) (@37°C) | Physical constant |
| $\Sigma A_c$ | 4.5 mM | Cellular adenine nucleotides concentration | Adjusted |
| $\Sigma N_m$ | 2.2 mM | Free pyridine nucleotides concentration in the mitochondrial matrix | (1) |
| $\Sigma N_c$ | 2.0 mM | Free pyridine nucleotides concentration in the cytoplasm (Adjustable) | (1) |
| $k_{gpd}$ | 0.01/s | Consumption rate of G3P | (1) |
| $k_{pyr}$ | 0.01/s | Consumption rate of pyruvate | Added for stability in ETC inhibition |
| $k_{NADHm}$ | 0.1/s | Consumption rate of mitochondrial NADH | (1) |
| $k_{NADHc}$ | 0.1/s | Consumption rate of cytosolic NADH | (1) |
| $k_{ATP}$ | 0.04/s | Basal consumption rate of ATP | (1) |
| $k_{ATPCa}$ | 90/mM/s | Consumption rate of ATP activated by calcium | (1) |

#### Conservation relationships

$$\begin{aligned} \Sigma N_m &= [NAD^+]_m + [NADH]_m \\ \Sigma N_c &= [NAD^+]_c + [NADH]_c \\ \Sigma A_c &= [ATP]_c + [ADP]_c + [AMP]_c \\ 1 &= X_1 + 2X_2 + 3X_3 \end{aligned}$$

#### Adenylate kinase

$$J_{ADK} \quad = k_f\left([ADP]_c^2 - [ATP]_c[AMP]_c/K_{eq}^{AK}\right)$$

| Parameter | Value | Description | Ref. |
|---|---|---|---|
| $k_f$ | 1000 mM/s | Forward (AMP-forming) rate constant of adenylate kinase. The parameter was set arbitrarily large to ensure the equilibrium of ATP-ADP-AMP. | Estimated |
| $K_{eq}^{AK}$ | 0.931 | The equilibrium constant of adenylate kinase (AMP-forming). | (2) |

#### Glucokinase (GK)

$$J_{glu} = V_m \frac{[ATP]_c}{[ATP]_c + K_{ATP}} \frac{[Glc]^n}{[Glc]^n + K_{Glc}^n}$$

| Parameter | Value | Description | Ref. |
|---|---|---|---|
| $V_m$ | 0.011 mM/s | Max rate of glucokinase | (1) |
| $K_{ATP}$ | 0.5 mM | Michaelis constant for ATP | (1) |
| $K_{Glc}$ | 7 mM | Michaelis constant for glucose | (1) |
| n | 1.7 | Cooperativity for glucose | (1) |

#### Glyceraldehyde 3-phosphate dehydrogenase (GPD)

$$J_{gpd} = V_m \frac{[G3P]}{[G3P] + K_{G3P}} \frac{[NAD^+]_c}{[NAD^+]_c + K_{NAD}[NADH]_c}$$

| Parameter | Value | Description | Ref. |
|---|---|---|---|
| $V_m$ | 0.5 mM/s | Max rate of GPD (Adjustable) | (1) |
| $K_{G3P}$ | 0.2 mM | Michaelis constant for G3P | (1) |

| | | | |
|---|---|---|---|
| $K_{NAD}$ | 0.09 | Activation constant for cytosolic NAD/NADH ratio | (1) |

#### Lactate production by lactate dehydrogenase (LDH)

$$J_{LDH} = V_m \frac{[Pyr]}{[Pyr] + K_{Pyr}} \frac{[NADH]_c}{[NADH]_c + K_{NADH}[NAD^+]_c}$$

| Parameter | Value | Description | Ref. |
|---|---|---|---|
| $V_m$ | 1.2 mM/s | Max rate of LDH (Adjustable) | (1) |
| $K_{Pyr}$ | 0.0475 mM | Michaelis constant for pyruvate | (1) |
| $K_{NADH}$ | 1 | Activation constant for cytosolic NADH/NAD ratio | (1) |

#### Steady-state cytosolic calcium levels

$$[Ca^{2+}]_c = [Ca^{2+}]_R + k_A^{Ca} \frac{([ATP]_c)^n}{([ATP]_c)^n + (K_{ATP}[ADP]_c)^n}$$

| Parameter | Value | Description | Ref. |
|---|---|---|---|
| $[Ca^{2+}]_R$ | 90 nM | Resting cytoplasmic calcium concentration | (1) |
| $k_A^{Ca}$ | 250 nM | Maximal activated calcium concentration | (1) |
| $K_{ATP}$ | 25 | Activation constant for ATP/ADP ratio | (1) |
| $n$ | 4 | Cooperativity for ATP/ADP ratio | (1) |

#### Oscillating calcium levels

Independent oscillating calcium levels were only used in the section: oscillating calcium on mitochondrial bioenergetics and dynamics.

$$\begin{aligned} [Ca^{2+}]_c &= [Ca^{2+}]_R + k_A^{Ca}(Axe^{1-Ax})^B \\ x &= \frac{t}{T} - \lfloor \frac{t}{T} \rfloor \end{aligned}$$

| Parameter | Value | Description | Ref. |
|---|---|---|---|
| $[Ca^{2+}]_R$ | 90 nM | Resting cytoplasmic calcium concentration | (1) |
| $k_A^{Ca}$ | 250 nM | Maximal activated calcium concentration | (1) |
| $A$ | 5 | Asymmetric factor | Estimated from (3) |
| $B$ | 4 | Steepness factor | Estimated from (3) |

| | | | |
|---|---|---|---|
| $T$ | 2 minute | Period of calcium oscillations. | Estimated from (3) |

#### Pyruvate dehydrogenase (PDH)

We assume that pyruvate diffuses freely and fast across the inner mitochondrial membrane (IMM). Therefore, pyruvate is the same concentration in the cytosol and the mitochondrial matrix.

$$
\begin{aligned}
J_{PDH} &= V_m \frac{[Pyr]}{[Pyr] + K_{Pyr}} \frac{[NAD^+]_m}{[NAD^+]_m + K_{NAD}} \frac{(1+C)^2}{(1+C)^2(1+u_2) + u_2 u_1} \\
C &= [Ca^{2+}]_m / K_{Ca}
\end{aligned}
$$

| **Parameter** | **Value** | **Description** | **Ref.** |
|---|---|---|---|
| $V_m$ | 0.3 mM/s | Max rate of PDH | (1) |
| $K_{Pyr}$ | 0.0475 mM | Michaelis constant for pyruvate | (1) |
| $K_{NAD}$ | 81 | Activation constant for mitochondrial NAD/NADH ratio | (1) |
| $K_{Ca}$ | 50 nM | Activation constant for mitochondrial Ca | (1) |
| $u_1$ | 1.5 | Factor for calcium activation | (1) |
| $u_2$ | 1.1 | Factor for calcium activation | (1) |

#### Electron transport chain (ETC)

$$
J_{hr} = V_m \frac{[NADH]_m}{[NADH]_m + K_{NADH}} \frac{1 + k_A \Delta\Psi_m}{1 + k_B \Delta\Psi_m} F_{O_2}
$$

| **Parameter** | **Value** | **Description** | **Ref.** |
|---|---|---|---|
| $V_m$ | 22 mM/s | Max rate of ETC | (1) |
| $K_{NADH}$ | 3 mM | Michaelis constant for NADH | (1) |
| $k_A$ | -4.92 /V | thermodynamic potential factor | (1) |
| $k_B$ | -4.43 /V | thermodynamic potential factor | (1) |
| $F_{O_2}$ | 1 | Oxygen availability | (1) |

#### F1Fo ATPase (ATP synthase)

In this model, ATP synthase was lumped with the adenine nucleotide translocator (ANT). The reaction rate depended on cytosolic ADP levels and mitochondrial membrane potential.

$$
\begin{aligned}
J_{hf} &= V_m f_{ADP} f_{\Psi} f_{Ca} \\
J_{ANT} &= J_{hf}/H_{ATP} \\
f_{ADP} &= \frac{[MgADP]_c^{n_A}}{[MgADP]_c^{n_A} + K_{ADP}^{n_A}} \\
f_{\Psi} &= \frac{\Delta\Psi_m^{n_\Psi}}{\Delta\Psi_m^{n_\Psi} + K_{\Psi}^{n_\Psi}} \\
f_{Ca} &= 1 - \exp(-[Ca^{2+}]_m/K_{Ca}) \\
[\mathrm{MgADP}]_c &= 0.055[ADP]_c
\end{aligned}
$$

| Parameter | Value | Description | Ref. |
|---|---|---|---|
| $V_m$ | 8 mM/s | Max rate of ATP synthase (Adjustable) | (1) |
| $K_{ADP}$ | 20 µM | Apparent Michaelis constant for cytosolic MgADP | (1) |
| $n_A$ | 2 | Cooperativity for MgADP | (1) |
| $n_\Psi$ | 8 | Cooperativity for mitochondrial potential | (1) |
| $K_\Psi$ | 131.4 mV | Mid-activity constant for mitochondrial potential | (1) |
| $K_{Ca}$ | 0.165 µM | Activation constant for mitochondrial calcium | (1) |
| $H_{ATP}$ | 3 | H:ATP ratio | Adjusted |

#### Proton leak

The basal leak approaches about 20% of the electron transport rate at $\Delta\Psi_m$ of 160 mV. (1)

$$J_{hl} = P_H \exp(k_{lp} \Delta\Psi_m)$$

| Parameter | Value | Description | Ref. |
|---|---|---|---|
| $P_H$ | 0.0024 mM/s | Proton leak coefficient | (1) |
| $k_{lp}$ | 30.5/V | Mitochondrial membrane potential coefficient | (1) |

#### NADH shuttles

$$J_{TNADH} = T_{NADH} \frac{[NADH]_c}{[NADH]_c + [NAD^+]_c K_c} \frac{[NAD^+]_m}{[NAD^+]_m + [NADH]_m K_m}$$

| Parameter | Value | Description | Ref. |
|---|---|---|---|
| $T_{NADH}$ | 0.05 mM/s | NADH transport rate | (1) |
| $K_c$ | 0.002 | Affinity coefficients for cytoplasmic NADH/NAD | (1) |
| $K_m$ | 16.78 | Affinity coefficients for mitochondrial NAD/NADH | (1) |

#### Mitochondrial calcium uniporter (MCU)

$$\begin{aligned} J_{uni} &= P_{Ca} \frac{\delta}{e^{\delta}-1} \left( e^{\delta} \alpha_i [Ca^{2+}]_c - \alpha_m [Ca^{2+}]_m \right) \\ \delta &= Z_{Ca} \Delta\Psi_m / V_T \end{aligned}$$

| Parameter | Value | Description | Ref. |
|---|---|---|---|
| $P_{Ca}$ | 4 / s | Permeability of calcium | (1) |
| $Z_{Ca}$ | 2 | Valence of calcium | (1) |
| $\alpha_i$ | 0.341 | Activity of cytoplasmic calcium | (1) |
| $\alpha_m$ | 0.2 | Activity of mitochondrial calcium | (1) |

#### Mitochondrial Sodium-Calcium exchanger (NCLX)

We used the electron-neutral descriptor of NCLX since this model generated a smooth and monotonous increment of mitochondrial calcium levels upon increasing glucose levels.

$$\begin{aligned} J_{NCLX} &= V_m (AB - PQ)/D \\ D &= 1 + A + B + P + Q + AB + PQ \\ A &= ([Na^{+}]_c / K_{Na})^2 \\ B &= [Ca^{2+}]_m / K_{Ca} \\ P &= ([Na^{+}]_m / K_{Na})^2 \\ Q &= [Ca^{2+}]_c / K_{Ca} \end{aligned}$$

| Parameter | Value | Description | Ref. |
|---|---|---|---|
| $V_m$ | 0.075 mM/s | Max rate of NCLX | (1), adjusted |
| $K_{Ca}$ | 8 µM | Dissociation constant of Ca | (1) |
| $K_{Na}$ | 8.2 mM | Dissociation constant of Na | (1) |

#### Mitochondrial Dynamics

$$\begin{aligned} k_{fuse,1} &= k_0^{fuse} J_{ANT} / J_{hl} \\ k_{fiss,1} &= k_0^{fiss} \\ k_{fuse,2} &= 0.5 k_{fuse,1} \\ k_{fiss,2} &= 1.5 k_{fiss,1} \end{aligned}$$

| Parameter | Value | Description | Ref. |
|---|---|---|---|
| $k_0^{fiss}$ | 0.05 /mM | Conversion factor for fusion rate | (4) |

| | | | |
|---|---|---|---|
| $k_0^{fuse}$ | 0.05 /mM | Conversion factor for fission rate | Estimated |

#### Ordinary differential equations

$$
\begin{aligned}
\frac{d}{dt}[G3P] &= \frac{1}{V_i}\left(2J_{glu} - J_{GPD}\right) - k_{g3p}[G3P] \\
\frac{d}{dt}[Pyr] &= \frac{1}{V_i + V_{mtx}}(J_{GPD} - J_{LDH} - J_{PDH}) - k_{pyr}[Pyr] \\
\frac{d}{dt}[NADH]_c &= \frac{1}{V_i}(J_{GPD} - J_{LDH} - J_{NADHT}) - k_{nadhc}[NADH]_c \\
\frac{d}{dt}[NADH]_m &= \frac{1}{V_{mtx}}(J_{NADHT} + 4.6J_{PDH} - 0.1J_{hr}) - k_{nadhm}[NADH]_m \\
\frac{d}{dt}\Delta\Psi_m &= \frac{1}{C_{mito}}\left(J_{hr} - J_{hf} - J_{ANT} - J_{hl} - 2J_{uni}\right) \\
\frac{d}{dt}[Ca^{2+}]_m &= \frac{f_m}{V_{mtx}}(J_{uni} - J_{NCLX}) \\
\frac{d}{dt}[ATP]_c &= \frac{1}{V_i}(J_{ANT} - 2J_{Glu} + 2J_{GPD} + J_{AdK}) - (k_{ATP} + k_{CaATP}[Ca^{2+}]_c)[ATP]_c \\
\frac{d}{dt}[ADP]_c &= -\frac{d}{dt}[ATP]_c - \frac{J_{AdK}}{V_i} \\
\frac{d}{dt}X_2 &= k_{fuse,1}X_1^2 - \left(k_{fiss,1} + k_{fuse,2}X_1\right)X_2 + k_{fiss,2}X_3 \\
\frac{d}{dt}X_3 &= k_{fuse,2}X_1X_2 - k_{fiss,2}X_3
\end{aligned}
$$

#### Initial conditions

| State variable | Value | Description |
|---|---|---|
| $[G3P]$ | 2.8μM | Glyceraldehyde-3-phosphate |
| $[Pyr]$ | 8.5μM | Pyruvate |
| $[NADH]_c$ | 1μM | Cytosolic NADH |
| $[NADH]_m$ | 60μM | Mitochondrial NADH |
| $[ATP]_c$ | 4mM | Cytosolic ATP concentration |
| $[ADP]_c$ | 0.5mM | Cytosolic ADP concentration |
| $[Ca^{2+}]_m$ | 0.250μM | Mitochondrial calcium concentration |
| $\Delta\Psi_m$ | 100mV | Mitochondrial membrane potential |
| $X_2$ | 0.0 | The population of degree-2 mitochondrial nodes |
| $X_3$ | 0.0 | The population of degree-3 mitochondrial nodes |
